# SARS-CoV-2 Omicron potently neutralized by a novel antibody with unique Spike binding properties

**DOI:** 10.1101/2022.03.18.484873

**Authors:** Craig Fenwick, Priscilla Turelli, Dongchun Ni, Laurent Perez, Kelvin Lau, Erica Lana, Céline Pellaton, Charlène Raclot, Line Esteves-Leuenberger, Jérémy Campos, Alex Farina, Flurin Fiscalini, Cécile Herate, Romain Marlin, Rana Abdelnabi, Caroline S. Foo, Johan Neyts, Pieter Leyssen, Roger LeGrand, Yves Lévy, Florence Pojer, Henning Stahlberg, Didier Trono, Giuseppe Pantaleo

**Affiliations:** Service of Immunology and Allergy, Department of Medicine, Lausanne University Hospital and University of Lausanne, Lausanne, Switzerland; School of Life Sciences, Ecole Polytechnique Fédérale de Lausanne, Lausanne, Switzerland; School of Basic Sciences, Ecole Polytechnique Fédérale de Lausanne and Faculty of Biology and Medicine, UNIL, Lausanne, Switzerland; KU Leuven Department of Microbiology, Immunology and Transplantation, Rega Institute for Medical Research, Laboratory of Virology and Chemotherapy, B-3000 Leuven, Belgium; CEA, Université Paris Sud 11, INSERM U1184, Center for Immunology of Viral Infections and Autoimmune Diseases, IDMIT Department, IBFJ, Fontenay-aux-Roses, France; VRI, Université Paris-Est Créteil, Faculté de Médicine, INSERM U955, 94010 Créteil, France; Inserm U955, Equipe 16, Créteil, France; AP-HP, Hôpital Henri-Mondor Albert-Chenevier, Service d’Immunologie Clinique et Maladies Infectieuses, Créteil, France; Swiss Vaccine Research Institute, Lausanne University Hospital and University of Lausanne, Switzerland

**Keywords:** SARS-CoV-2, neutralizing antibodies, Variants of concern, Omicron

## Abstract

The SARS-CoV-2 Omicron variant exhibits very high levels of transmission, pronounced resistance to authorized therapeutic human monoclonal antibodies and reduced sensitivity to vaccine-induced immunity. Here we describe P2G3, a human monoclonal antibody (mAb) isolated from a previously infected and vaccinated donor, which displays picomolar-range neutralizing activity against Omicron BA.1, BA.1.1, BA.2 and all other current variants, and is thus markedly more potent than all authorized or clinically advanced anti-SARS-CoV-2 mAbs. Structural characterization of P2G3 Fab in complex with the Omicron Spike demonstrates unique binding properties to both down and up spike trimer conformations at an epitope that partially overlaps with the receptor-binding domain (RBD), yet is distinct from those bound by all other characterized mAbs. This distinct epitope and angle of attack allows P2G3 to overcome all the Omicron mutations abolishing or impairing neutralization by other anti-SARS-COV-2 mAbs, and P2G3 accordingly confers complete prophylactic protection in the SARS-CoV-2 Omicron monkey challenge model. Finally, although we could isolate *in vitro* SARS-CoV2 mutants escaping neutralization by P2G3 or by P5C3, a previously described broadly active Class 1 mAb, we found these viruses to be lowly infectious and their key mutations extremely rare in the wild, and we could demonstrate that P2G3/P5C3 efficiently cross-neutralized one another’s escapees. We conclude that this combination of mAbs has great prospects in both the prophylactic and therapeutic settings to protect from Omicron and other VOCs.

---

SARS-CoV-2 has to this day been responsible for >340 million confirmed infections and >5.5 million fatalities ^1^. Its unabated propagation has progressively led to the emergence of variants of concern (VOC) with enhanced transmission and resistance to immune responses. The scene has been most recently dominated by VOCs harbouring a high number of mutations compared to the original SARS-CoV-2 strain, with Delta (B.1.617.2) and its 11 to 15 Spike mutations now supplanted by the highly infectious Omicron (B.1.1.529.1), which contains up to 37 amino acid mutations in this viral protein ^2,3^. Fifteen of Omicron Spike substitutions reside within the RBD, the region targeted by most neutralizing antibodies whether induced by infection or current vaccines, all derived from the original 2019-nCoV Wuhan strain ^4–8^. Omicron also resists neutralization by most anti-SARS-CoV-2 mAbs reported so far ^9–15^ and is circulating as several sub-variants including BA.1.1, BA.2 (B.1.1.529.2) and BA.3 (B.1.1.529.3) ^2^, creating an urgent unmet medical need for both prophylaxis and therapeutics.

## Results

### Identification of P2G3, a highly potent SARS-CoV-2 neutralizing antibody

We screened for the presence of anti-Spike antibodies in serum samples from a cohort of >100 donors and focused on one post-infected donor that received two doses of the mRNA-1273 vaccine and displayed among the highest serum antibody levels with excellent breadth against a panel of SARS-CoV-2 variants in a trimeric Spike-ACE2 surrogate neutralization assay ^16^. Screening of B cell clone supernatants for high affinity Spike binding led us to prioritize six clones for mAb production via expression of paired heavy and light chains in ExpiCHO cells. During initial profiling of these purified mAbs, P2G3 exhibited the strongest binding affinity for the original 2019-nCoV Spike trimer and a panel of Spike proteins encoding mutations found in Alpha, Beta, Gamma and Delta VOCs (IC_50s_ of 0.006-0.010 μg/ml) (**Extended Data Fig. 1a)**. Cross-competitive Spike RBD binding studies performed with a panel of authorized or clinically advanced anti-SARS-CoV-2 mAbs (REGN10933 and REGN10987 from Regeneron ^17^, AZD8895 and AZD1061 from AstraZeneca ^18^, ADG-2 from Adagio ^19^, S309/Sotrovimab from Vir/GSK ^20^) and mAbs previously described by our group ^21^, demonstrate that P2G3 binds an unique albeit overlapping epitope with those recognized by both AZD1061 and S309/Sotrovimab, the latter of which acts by a mechanism distinct from blocking the RBD/ACE2 interaction^20^ (**Extended Data Fig. 1b)**. Importantly, our potent and broadly active Class 1 mAb, P5C3, bound RBD non-competitively with P2G3, prompting us to profile these mAbs both alone and in combination for subsequent studies.

Using a biochemical trimeric Spike-ACE2 surrogate neutralization assay ^16^, we further determined that P2G3 and P5C3 had the most potent and broad activity in blocking ACE2 binding to high quality structural grade Spike trimers from all past VOCs compared to our panel of benchmark mAbs (**Extended Data Fig. 1 c-d**). P2G3 and P5C3 also display the most potent activity in assays performed with the Spike protein from Omicron BA.1, BA.1.1 that includes the R346K mutation and BA.2. (**Fig. 1a and 1b**). P2G3 alone gave comparable activities to the P2G3/P5C3 combination, with the cocktail showing improved IC_80_ values of 6.1-, 47.2- and 3.4-fold versus the AZD1061/AZD8895 mix, 9.9-, 12.4 and 340-fold versus ADG-2 and 375-, 337- and 32-fold versus the REGN10933/ REGN10987 mix in assays performed with Omicron BA.1, BA.1.1 and BA.2, respectively. Interestingly, while the P2G3/P5C3 combination reached ~100% of inhibition in blocking ACE2 binding to Omicron Spike proteins at 1 μg/ml total mAbs, P2G3 alone only blocked 50-72% of ACE2 binding at 20 μg/ml and P5C3 alone reached a ~100% inhibition at 20 μg/ml.

**Figure 1:**
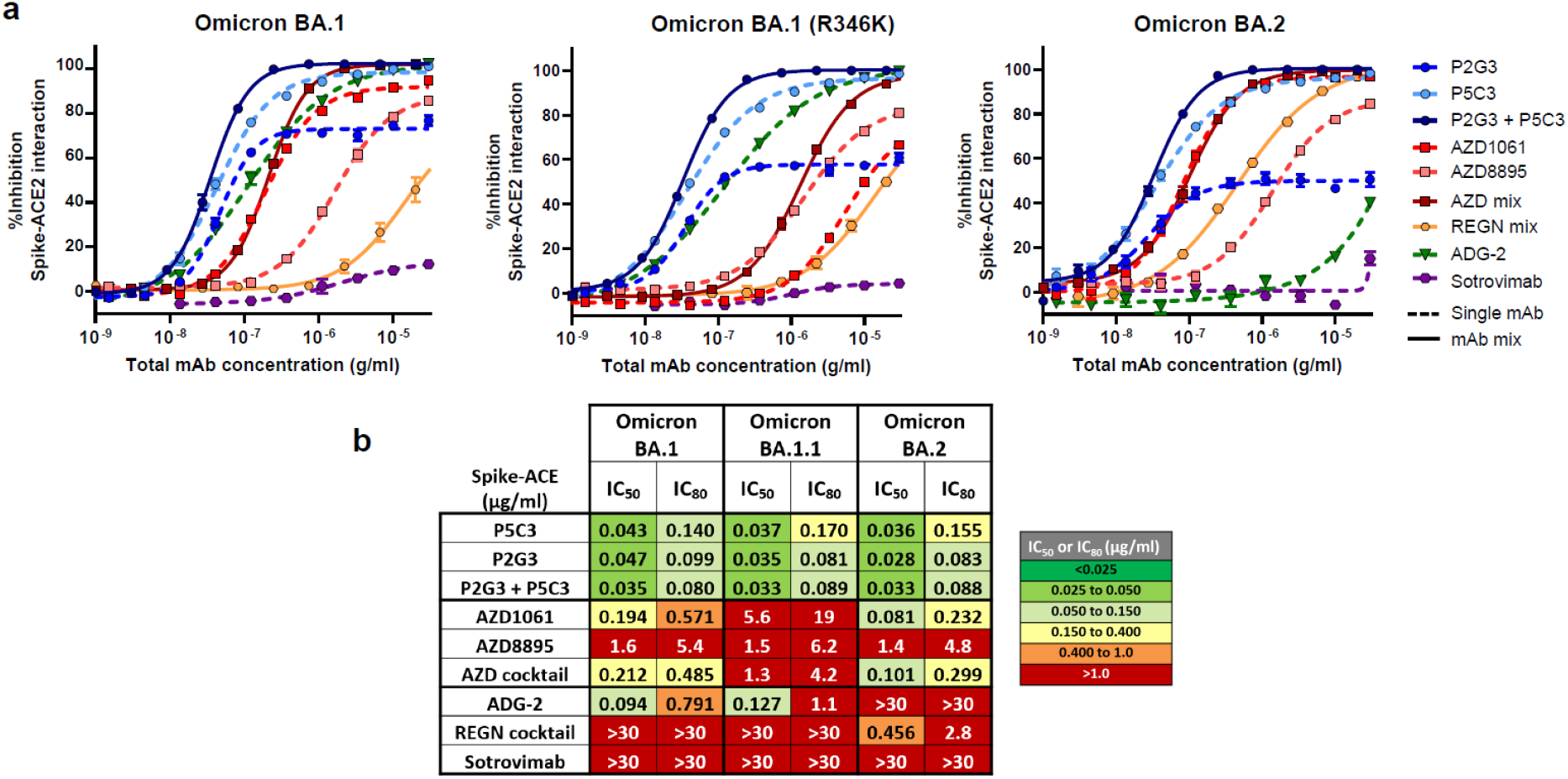
Identification of P2G3, a human mAb with potent activity in a Spike-ACE2 surrogate neutralization assay performed with Omicron BA.1, BA.1.1 and BA.2 trimeric Spike proteins. **a)** Blocking activity of individual and combinations of anti-Spike mAbs in a biochemical Spike-ACE2 surrogate neutralization assay using Omicron BA.1, BA.1.1 encoding the R346K mutation and BA.2 variant Spike trimer proteins. **b)** Comparative Spike-ACE2 blocking activity of P2G3, P5C3 and the P2G3/P5C3 mix compared to a panel of benchmark anti-SARS-CoV-2 mAbs. P2G3 and P5C3 used in these studies and throughout the manuscript contain the LS mutation in the Fc domain (M428L/N434S), previously demonstrated to confer an extended half-life *in vivo*. Data presented is representative of 2-4 independent experiments with each concentration response tested in duplicate. Mean values ± SEM are shown in a).

We next compared P2G3 alone or in combination with P5C3 with our panel of current clinically authorized and/or clinically advanced mAbs in pseudovirus neutralization assays. P2G3 had potent neutralizing activity against lentiviruses pseudotyped with Spike from initial 2019-nCoV (D614G), Alpha, Beta and Delta VOCs (IC_80_ value of 0.022, 0.051, 0.038 and 0.035 μg/ml, respectively). Most importantly, P2G3 strongly neutralized the Omicron BA.1 Spike pseudovirus with an IC_80_ value of 0.038 μg/ml (**Fig. 2a**) and thus showed no loss of activity as compared to the other VOCs. In side-by-side comparisons, P2G3 was >42-fold more potent than ADG-2, AZD1061, AZD8895, REGN10933 and REGN10987 mAbs and 19-fold more potent than Sotrovimab at neutralizing Omicron BA.1 Spike pseudotyped lentiviral particles (**Fig. 2b**). Second most potent was P5C3, with an IC_80_ value of 0.223 μg/ml, and the P2G3/P5C3 combination revealed a minor-enhanced activity over P2G3 alone in this assay, with an IC_80_ value of 0.024 μg/ml for the total concentration of the two mAbs. Furthermore, P2G3 and P5C3 maintained full neutralizing activity against the ancestral D614G and Omicron BA.1.1 encoding the R346K Spike pseudovirus (**Extended Data Fig. 2a and b**), a mutation present in ~10% of Omicron variant sequences in the GISAID ^11^.

**Figure 2:**
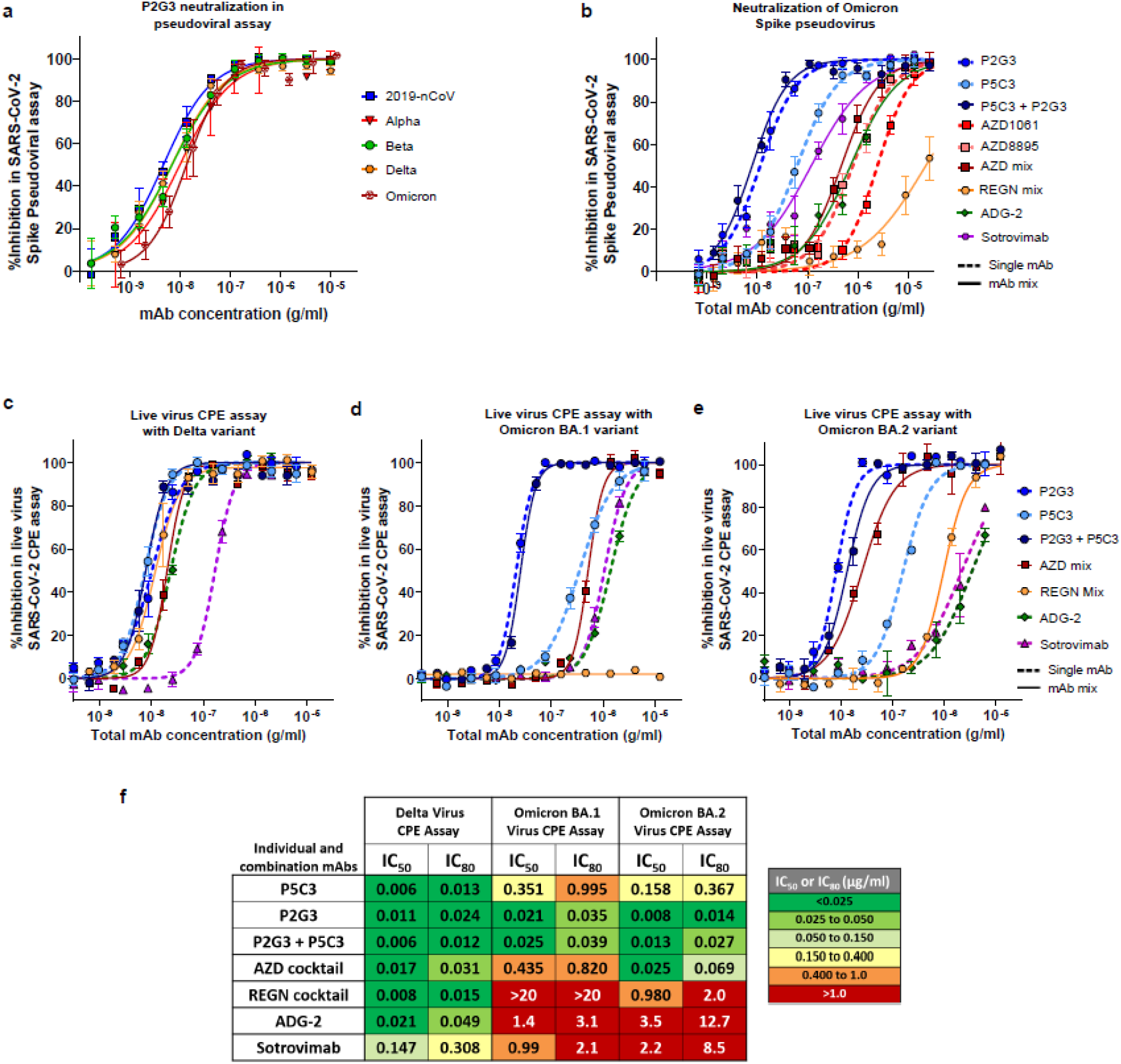
P2G3 demonstrates potent and broad neutralization of Spike-coated pseudoviruses and live virus SARS-CoV-2 VOCs. **a**) Neutralization of lentiviral particles pseudotyped with SARS-CoV-2 Spike expressing variants of concern in a 293T-ACE2 infection assay. All Spike proteins contained the D614G substitution that became dominant early in the pandemic. **b**) Neutralization of lentiviral particles pseudotyped with the Omicron variant Spike. Antibody cocktails representing a 1:1 mix of each mAb to give the indicated total mAb concentration. **c-e)** Neutralization activity of P2G3 performed in a live SARS-CoV-2 infectious virus cytopathic effect assay (CPE). The indicated SARS-CoV-2 variants were used to infect Vero E6 *in vitro* in the absence and presence of concentration response of the indicated mAb. Viral inhibition curves are shown for the c) Delta, d) Omicron BA.1 and e) BA.2 variants for individual and mAb combinations. **f**) Heatmap showing IC_50_ and IC_80_ neutralization potencies for the indicated mAbs in the live virus CPE assays. Results shown in a-d are the average of two to three independent experiments with each concentration response tested in duplicate or triplicate. Results in f) are from an individual experiment with each concentration tested in duplicate. Mean values ± SEM are shown.

We next profiled P2G3 using the initial D614G strain and all current VOCs in a live virus cytopathic effect assay. P2G3 demonstrated broad and potent neutralizing activity with IC_80_ values of 0.028, 0.010, 0.017, 0.021, 0.024 and 0.035 μg/ml against the 2019-nCoV (D614G) strain, Alpha, Beta, Gamma, Delta and Omicron variants, respectively (IC_80_ ranging from 67-233 pM) (**Extended Data Fig. 2c**). We went on to compare P2G3 with other mAbs alone or in combinations for their ability to block the Delta and currently prevalent Omicron BA.1 and BA.2 SARS-CoV-2 variants (**Fig. 2c-2f**). All tested mAbs displayed good activity against the Delta variant, although Sotrovimab was ~6 times less potent than ADG-2 and >10 times less potent than P2G3, P5C3 and both the AZD and REGN cocktails against this virus. However and most notably, P2G3 was by far the most active mAb against Omicron BA.1 with an IC_80_ value of 0.035 μg/ml, that is ~23-fold, 60-fold and 88-fold more potent than those of AZD1061/AZD8895, Sotrovimab and ADG-2, respectively, the REGN combination being completely ineffective against this variant (**Fig. 2d and 2f**). Of note, while P5C3 alone displayed an IC_80_ against Omicron in the range of that of the AZD cocktail, the P2G3/P5C3 combination showed high neutralization activity comparable to P2G3 alone with an IC_80_ of 0.039 μg/ml total mAb. P2G3 was again the most potent mAb in neutralizing the Omicron BA.2 variant with an IC_80_ values of 0.014 μg/ml, that is 4.9-fold, 14-fold, 907-fold and 607-fold more potent than AZD1061/AZD8895, REGN10933/REGN10987, ADG-2 and Sotrovimab, respectively (**Fig. 2e-2f**). Importantly, although the Spike-ACE2 surrogate neutralization assay correlates well with cell-based neutralization assays ^16^, P2G3 only blocked ~50-72% of ACE2 binding to Omicron Spike proteins (**Fig. 1a**) despite P2G3 reaching 100% of the maximum signal in a direct binding assay (**Extended Data Fig. 1e**). This suggests that at least with the Omicron variant, virus inhibition by P2G3 may not be solely mediated through the Spike-ACE2 interaction, but rather a mechanism analogous with that of S309/Sotrovimab, reported to be through induction of Spike trimer cross-linking, steric hindrance, aggregation of virions ^20^ and/or inhibiting viral membrane attachment through C-type lectin receptors ^22^.

Given the potential importance of Fc-mediated antibody effector functions in more efficient virus control and clearance ^23–25^, we investigated P2G3 activities in antibody dependent cellular cytotoxicity (ADCC) and antibody dependent cellular phagocytosis (ADCP) assays. ADCC enables the targeted killing of cells that display SARS-CoV-2 Spike protein at the membrane surface and our *in vitro* assay uses CEM-NKR cells stably expressing 2019-nCoV Spike cultured with primary effector cells from healthy donors in the presence and absence of anti-Spike mAbs. P2G3 mAb exhibited a robust ADCC activity that was superior in killing Spike positive cells compared to all other anti-Spike mAbs tested (**Extended Data Fig. 3a**). Although these studies were not performed with an Omicron Spike stable cell line, it is generally accepted that ADCC functional activity is maintained against the different SARS-CoV-2 VOCs for mAbs that conserve Spike variant binding ^26^. We next evaluated ADCP activity using 2019-nCoV or Omicron BA.1 Spike trimer coated fluorescent beads mixed with different concentrations of P2G3 and/or P5C3 then incubated with U937 effector cells. This monocyte cell line expresses high levels of Fc-gamma receptors capable of inducing phagocytosis of opsonized viruses or beads coated with the Spike antigen as in our assay (**Extended Data Fig. 3b**). Used alone, P2G3 and P5C3 mAbs showed ADCP IC_80_ activities of 0.074 and 0.010 μg/ml, respectively with P5C3 showing ~7-fold greater potency. Conversely, using Omicrom Spike coated beads, P2G3 exhibits potent ADCP activity that was 3-fold improved relative to P5C3. In studies performed with both ancestral and Omicron Spike, the P2G3/P5C3 mix shows enhanced ADCP activities compared to mAbs used individually (**Extended Data Fig. 3c-d**). Of note, P2G3 and P5C3 used in these Fc-mediated functional activity assays and throughout the manuscript contain the LS mutation in the Fc domain (M428L/N434S), previously demonstrated to confer an extended half-life *in vivo* ^27^, a highly desirable feature for prophylactic use.

### P2G3 confers strong *in vivo* prophylactic protection from ancestral 2019-nCoV and Omicron SARS-CoV-2 infection

Having demonstrated the superior *in vitro* neutralizing activity of P2G3,we next evaluated the neutralizing potency of P2G3 *in vivo* in a prophylactic hamster challenge model of SARS-CoV-2 infection. Antibody dosed animals were challenged two days later with an intranasal inoculation of the original 2019-nCoV SARS-CoV-2 virus (**Extended Data Fig. 4 a**) and then four days later, hamster lung tissue was monitored for infectious virus and viral RNA. Infectious virus was undetectable in lungs from almost all P2G3 treated hamsters, with only 1 of 6 hamsters in the lowest dosed 0.5 mg/kg group showing reduced though detectable levels of infectious virus (**Extended Data Fig. 4 b**). Complete prophylactic protection was observed with P2G3 mAb plasma levels >6.2 μg/ml at the time of viral inoculation and P2G3 treatment groups showed a significant ~4-log reduction of genomic viral RNA levels (**Extended Data Fig. 4 c)**.

We next evaluated P2G3 mediate protection from SARS-CoV-2 Omicron BA.1 infection in a cynomolgus macaques pre-exposure challenge study. Monkeys were administered 10 mg/kg of P2G3 LS intravenously and challenged 72 hrs later via combined intranasal and intratracheal routes with 1×10^5^ TCID50 of SARS-CoVo-2 B.1.1.519 Omicron BA.1 virus (**Fig. 3a**). Following viral challenge, control animals showed similar genomic (g)RNA levels and kinetics with median peak viral loads (VL) of 6.9- and 6.6-log10 copies/ml gRNA at 2-3 days post challenge in tracheal swabs and bronchoalveolar lavage (BAL) samples, respectively (**Fig. 3b**). Nasopharyngeal swabs showed a higher-level variability in VL between control animals but still showed median peaks of 6.9-log10 copies/ml for gRNA. In comparison, the two P2G3 LS treated monkey had a strong median peak VL reduction of 3.8-, 2.5- and 3.9-log10 copies/ml gRNA for tracheal, nasopharyngeal and BAL samples, respectively.

**Figure 3.**
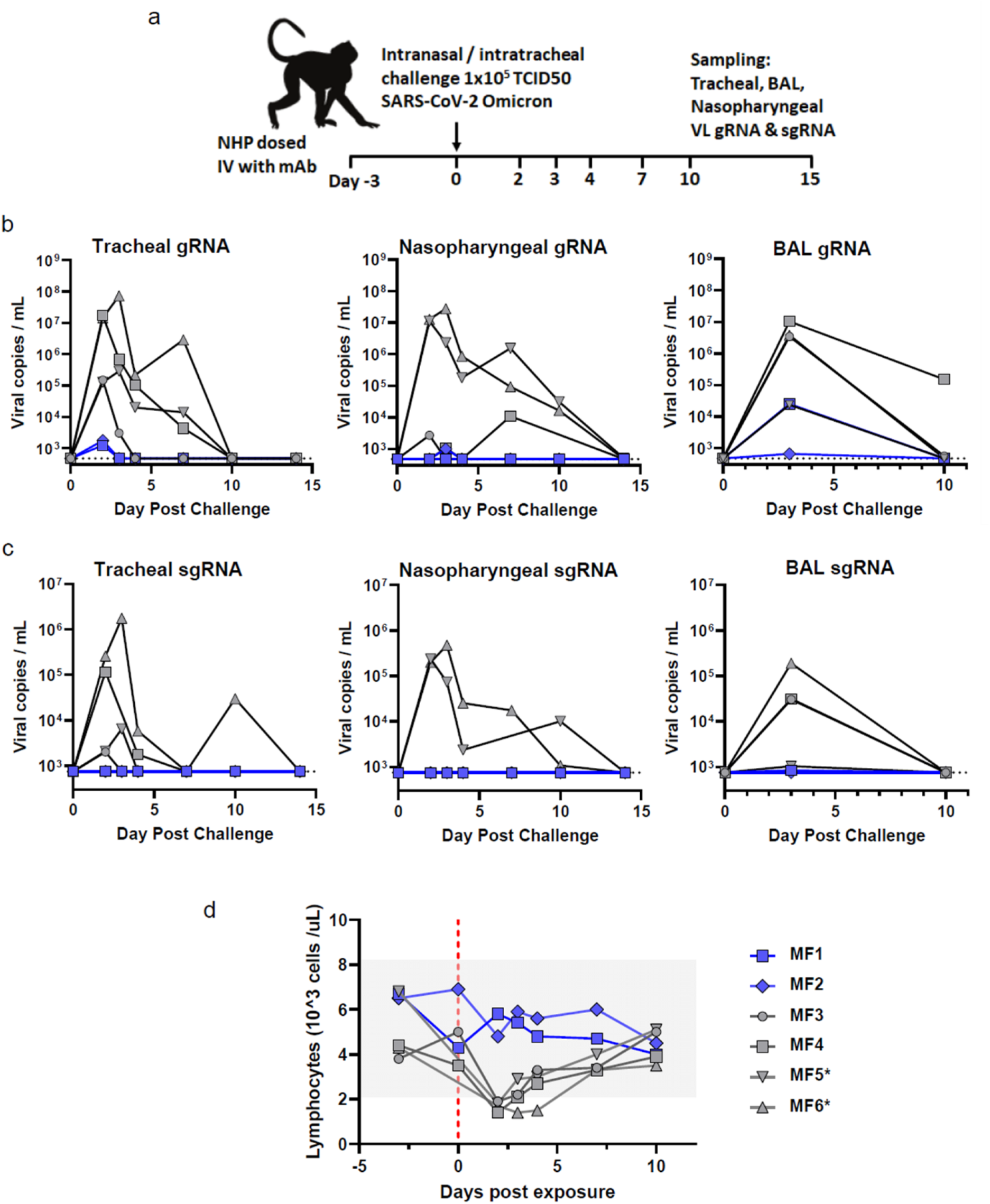
P2G3 LS confers potent *in vivo* efficacy in the non-human primate (NHP) challenge model for Omicron BA.1 SARS-CoV-2 infection. **a)** Overview of study design for the SARS-CoV-2 NHP challenge model. Animals MF1 and MF2 were administered intravenous 10 mg/kg of P2G3 and challenged three days later (Day 0) along with control animals MF3 and MF4 via intranasal and intratracheal inoculation of the Omicron BA.1 SARS-CoV-2 virus (1 x10^5^ TCID_50_). Tracheal swabs, nasopharyngeal swabs and bronchoalveolar lavages (BAL) performed during the course of the study were evaluated for viral copies per ml of genomic (g)RNA **b)** and subgenomic RNA **c)** with data plotted to include two historical control animals (MF5* and MF6*) infected with the same inoculum of Omicron virus. **d)** Flow cytometry analysis of blood samples from NHPs collected throughout the study shows strong lymphopenia in control animals following challenge with Omicron SARS-CoV-2 while P2G3 LS treated monkeys shows stable levels of lymphocytes. Dotted line indicates lower limit of detection at 2.68- and 2.87-log copies per ml for viral gRNA and sgRNA, respectively.

Active viral replication, as assessed by subgenomic (sg)RNA levels, peaked 2-3 days post-challenge with tracheal swabs and BAL showing median values of 4.8- and 4.5-log10 copies per ml, respectively and nasopharyngeal samples showing variable responses from undetectable (<2.9-log10) to 5.4-log10 copies/ml (**Fig. 3c**). P2G3 LS treated monkeys had sgRNA levels that were at or below the limit of detection, exhibiting 1.9-, 2.2- and 1.6-log10 reduced levels in tracheal, nasopharyngeal and BAL samples, respectively. Consistent with viral protection resulting in reduced detection of gRNA and sgRNA, P2G3 treated monkeys exhibited stable lymphocyte levels throughout the study, whereas strong lymphopenia, determined by lymphocyte levels below 2.1x 10^3^ cells/μl, was observed in all control animals challenged with the Omicron variant of SARS-COV-2 (**Fig. 3d**).

### Structural analysis of P2G3 and P5C3 Fab bound to Omicron Spike trimer

To decipher the molecular features underlying P2G3 and P5C3 potent neutralization of Omicron Spike, we performed single particle cryo-EM reconstruction of the Omicron Spike trimeric ectodomain ^12,28,29^ bound to both Fabs, at a 3.04 Å resolution (**Fig. 4a, Extended Data Fig. 5–6 and Extended Data Table 1**). We found the Fabs to bind simultaneously at distinct sites on the trimer with a majority of images revealing three P2G3 Fabs bound to either up- or down-RBD conformations and one P5C3 Fab bound to an up-RBD. The region of the Omicron RBD interacting with the Class 1 P5C3 mAb is identical to that previously described for the D614G Spike **(Extended Data Fig. 5–7**)^21^. P2G3 binds a surface area of around 700 Å^2^ as a Class 3 neutralizing mAb ^4^, recognizing an epitope on the SARS-CoV-2 RBD distinct from the receptor-binding motif, with the two mAbs covering together a surface of greater than 1200 Å^2^ (**Fig. 4b** and **Extended Data Fig. 7a, 8a**). To characterize the P2G3 paratope and epitope interface in detail, we performed local refinement of the P2G3 Fab-RBD interacting region and reached a resolution of 3.84 Å with well-defined density, allowing clear interpretation of sidechain positions (**Extended Data Fig. 6, 8–9** and **Extended Data Table 1)**. The P2G3 paratope is composed of four complementarity-determining region (CDR) loops binding at the back of the RBD. The interactions are mediated through electrostatic and hydrophobic contacts (**Fig. 4c-d** and **Extended Data Fig. 8b-c)** and involve sixteen residues of the RBD, mainly bound by the heavy chain of the P2G3 mAb. The 18-residue-long CDRH3 sits at the top of a loop that comprises residues 344–347, and also contacts the amino-acids at the limits of the 5-stranded β-sheet (residues 440–451), overall accounting for more than 60% of the buried surface area (431 Å^2^) (**Extended Data Fig. 8a-c**). The interactions between P2G3 and the Omicron RBD are conserved in both RBD-up and RBD-down states (**Extended Data Fig. 8c-d)**. CDRH2 extends the epitope by interacting with R346 that is engaged by residue W53 by a potential cation-pi interaction (**Fig. 4d and Extended Data Fig. 8e**), an interaction that is likely conserved with the R346K Spike substitution (**Extended Data Fig. 2a-b**). The only potential contact from the light chain derives from the CDRL1 Y32 forming a hydrophobic interaction with V445 of the RBD (**Extended Data Fig. 8c**). Moreover, P2G3 is only observed to contact RBD amino acid residues and the distance to the nearest atom of the glycan branch is ~10Å from P2G3. Importantly, the epitope defined by our structural studies rationalizes the potent neutralizing activity of P2G3 against the Omicron variant relative to other Class 3 mAbs. Omicron mutations S371L, N440K, G446S and the minor R346K sub-variant are all situated outside of, adjacent to or have little effect on recognition of the P2G3-binding epitope, whereas two or more of these mutations directly impinge on epitopes recognized by REGN10987, AZD1061 and S309/Sotrovimab (**Fig. 4e**). Furthermore, P2G3 displays a unique binding orientation on the RBD with its Fab angling away from most of these Omicron mutations (**Fig. 4f**). In modelling the observed angles of attack of various Class 3 mAbs, it is likely that REGN10987 only binds to the up-RBD form while AZD1061 binds one up- and one down-RBD form, with steric hindrance blocking the third RBD site on the Omicron Spike trimer (**Extended data Fig. 10**). In contrast, P2G3 and S309/Sotrovimab likely binds both the up and down forms of Omicron RBD without clashes (**Fig. 4g, Extended data Fig. 10c**), a characteristic that may contribute to the largely conserved and high potency of P2G3 across VOCs.

**Figure 4.**
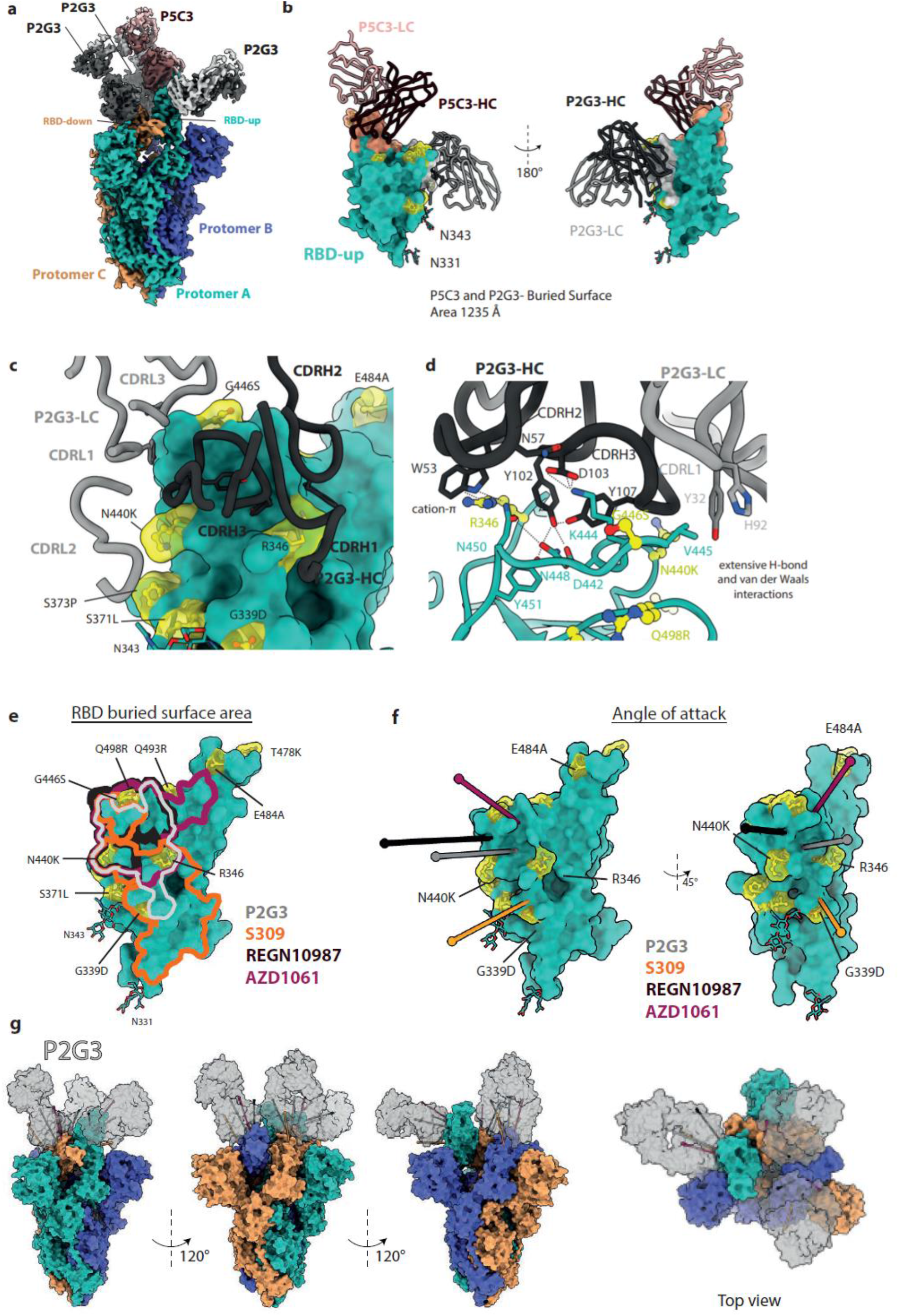
Two potent neutralizing antibodies P2G3 and P5C3 bind the full-length Omicron Spike. **a**) Cryo-EM composite density map of the full-length Omicron Spike bound to one P5C3 and three P2G3 Fab fragments. Spike protomers are colored in green, orange and blue, P5C3 Fabs in dark and light pink, P2G3 in black and grey. **b**) Surface representation of the RBD in the up configuration (green) bound to both P5C3 and P2G3 with heavy and light chains depicted as liquorice ribbons. The buried surface area formed by the Fabs are depicted on the RBD surface and coloured in grey for P2G3 and pink P5C3. The Omicron mutations are shown in yellow as balls-and-sticks and transparent surfaces. The N-linked glycans at asparagine 331 and 343 are shown as sticks. **c**) Zoomed-in view of the interacting region of P2G3 with CDR loops of the heavy and light chains specified. Omicron mutations are highlighted in yellow. **d**) Detailed analysis of the interactions between the Omicron RBD shown as ribbons (green) and the P2G3 Fab heavy and light chains shown as liquorice (black and grey). Residues at the interface are shown as sticks with potential interactions of interest as dashed lines. Omicron mutations are shown as balls-and-sticks in yellow. **e)** Structures of the several class 3 antibodies (Fabs) bound to RBDs were superimposed on the Omicron RBD. The buried surface area formed by the indicated Fabs are outlined on the RBD surface and coloured correspondingly (Fab-RBD structures AZD1061, PDB-7L7E; REGN10987, PDB-6XDG; S309, PDB-7BEP). The Omicron mutations are shown in yellow as balls-and-sticks and transparent surfaces. **f)** Fab binding angle of attack to the RBD is defined as the line connecting the centroid of the Fab to the centroid of the surface area of the RBD that the Fabs bury. Angle of attack of P2G3 compared to other class 3 antibodies viewed from multiple angles. RBD is in green, Omicron mutations in yellow. **g)** P2G3 binding to the full Omicron trimer was modelled by superimposing the Fabs on to the RBD of each protomer. The complex is shown from different sides and top view. P2G3 Fabs are able to bind all RBD-up and RBD-down conformations simultaneously.

### Cross-neutralization of P2G3 and P5C3 escape mutants

Monoclonal antibodies, as other classes of antivirals, are typically used in combination to prevent the emergence of resistant viruses. To gain insight into the predicted clinical value of our mAbs, we characterized the emergence of mutants escaping their blockade in tissue culture. For this, we grew SARS-CoV-2 Delta and Omicron BA.1 variants in the presence of sub-optimal neutralizing doses of either P2G3 or P5C3 for three passages to generate a heterogeneous viral population, before switching to stringent mAb concentrations in order to select *bona fide* escapees (**Fig. 5a**). Viral genome sequencing of these mAb-resistant mutants pointed to the importance of Spike substitutions G476D, F486S and N487K/D/S for escaping P5C3, and K444T for avoiding P2G3 neutralization. We thus tested the impact of these mutations on viral infectivity using lentivector pseudotypes. P5C3-escaping Spike proteins were significantly less infectious than the wild-type control in both the ancestral D614G and Delta backgrounds, correlating with a drop in affinity for the viral ACE2 receptor in an *in vitro* binding assay, whereas the P2G3-escaping K444T substitution had a milder effect in both assays **(Fig. 4b-c)**. Yet, an examination of the GISAID EpiCoV database revealed that mutations G476D, F486S or N487K/D/S found in P5C3 escapees are only exceptionally encountered in SARS-Cov2 isolates, representing together only 0.0087% of the 8’568’006 available sequences as of March 2022, and that the K444T mutation is equally rare (0.0024% of compiled sequences), strongly suggesting that the corresponding viruses have a markedly reduced fitness in the wild. Moreover, cross-neutralization studies demonstrated that P2G3 completely blocked the infectivity of P5C3-escaping Delta derivatives (**Fig. 4d**), and P5C3 and P2G3 efficiently cross-neutralized each other’s escape mutants in both the Delta and Omicron backgrounds (**Fig. 4e and Extended data Fig. 11**).

**Figure 5:**
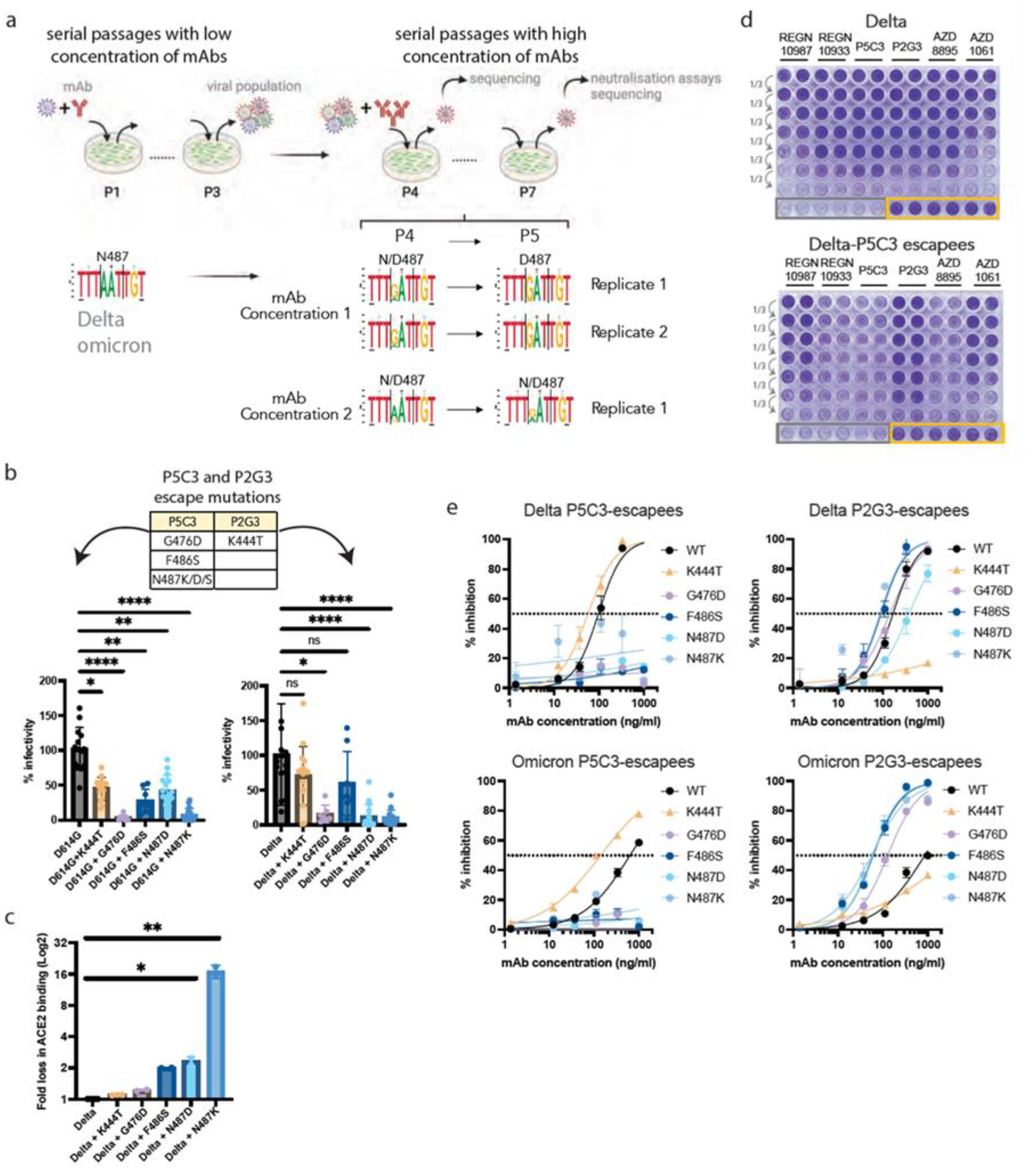
Identification and characterization of escape mutations to P2G3 and P5C3. **a)** Schematic representation of escapees selection. Delta and Omicron replicative isolates were used to infect VeroE6 cells (MOI of 0.2) each in duplicates in presence of suboptimal concentrations of antibodies. Supernatants were collected, diluted 40-fold and used to infect cells for two more passages in the same conditions (P1 to P3). Putative viral escapees were further selected by serial passages of 2-fold diluted supernatants pre-incubated with high concentrations of antibodies (three concentrations, each tested in duplicates). Viral RNA extracted from supernatants collected at each passage was deep-sequenced and P5 viral supernatant used for CPE-based neutralization assays. **b)** Mutations identified across escape selection experiments are indicated in the table. Lentivectors pseudotyped with Spikes mutated on the identified residues were produced in parallel (n=1 to 4 productions replicates). Lentiviral stocks were adjusted for p24 content and the same amount of each lentivector used to transduce 293T ACE2 cells (n=8 technical replicates for each transduction). Transduction efficiency was monitored by Luciferase activity in the transduced cells. **c)** The direct binding of ACE2 to mutated trimeric Spikes was monitored by Luminex-based binding assays. **d)** VeroE6 cells were infected in duplicates with normalized amounts of Delta- or Delta P5C3-escapees collected from escapees selection experiments and pre-incubated or not with 3-fold serial dilutions of mAbs as indicated. Cytopathic effect was monitored 2 days later with crystal violet staining of the live cells. Grey and yellow squares: respectively cells infected with viruses in absence of mAbs and non-infected cells. **e)** Luminex-based cross-neutralization assays were performed with either P5C3 or P2G3 antibodies on delta- or omicron-Spike derivatives. Kruskal-Wallis tests with Dunn’s multiple-comparison correction was performed to compare wild type and mutants in panels b and c; *p<0.05, **p<0.01, ***p<0.001, ****p<0.0001.

## Discussion

In sum, we report the discovery of P2G3, an anti-SARS-CoV-2 mAb with superior breadth and potency for neutralization of all VOCs, including the recently identified Omicron BA.1 and BA.2 variants. Structural and competitive binding studies demonstrate that P2G3 is a Class 3 mAb recognizing an epitope on the RBD different from those bound by all other therapeutic mAbs authorized or at an advanced stage of development^8,10,11^. Sequence analyses also reveal that the P2G3 HCDR3 presents only 55% identity with any of the 4897 anti-Spike HCDR3 described thus far. Cryo-EM studies further demonstrate that P2G3 can bind both the up- and down-RBD conformations of the Spike trimer, burying a large surface area of around 700 Å^2^ in a highly conserved region of RBD. Importantly, the binding epitope is largely non-overlapping with residues mutated in Omicron, a feature unique among almost all potent anti-SARS-CoV-2 mAbs reported so far ^11^. The S371L Omicron mutation alone remarkably reduces neutralization activity of multiple potent mAbs of different binding classes despite not being included in their footprint ^11^, perhaps because local conformational changes in the 370-375 loop impact the up- and down-RBD states and/or interfere with the critical gate opener N343 glycan positioning ^30^, yet this mutation is without effect on P2G3 neutralization. The unique angle of attack of P2G3 on the RBD domain, predicted to allow binding in both up- and down-RBD positions of the Spike trimer, may explain why this antibody remains potently active against the Omicron variant. Interestingly, we predict that S309/Sotrovimab is also sterically free to bind all RBDs within the Omicron trimer, yet we find this antibody to have a marked loss of activity against this variant. Although the epitope bound by P2G3 partly overlaps with the region recognized by S309-Sotrovimab, the latter mAb does not block the RBD/ACE2 interaction, and has been proposed to act by alternative mechanisms including the inhibition of cell adhesion through C-type lectins ^22^. Therefore, the improved binding affinity of P2G3 combined with the additional inhibitory effect on RBD/ACE2 interaction, explaining its 10 to 60-fold improved activity across all VOCs as compared to S309/Sotrovimab. Apart from the *in vitro* neutralization activity, P2G3 confers an excellent *in vivo* prophylactic protection in the SARS-COV-2 Omicron NHP challenge model. Our NHPs data also confirms studies in mice and hamsters indicating that the Omicron variant has reduced proliferation capacity in several animal models compared to other VOCs ^31,32^.

Despite the exceptional neutralization profile of P2G3 against currently circulating variants, development of resistance is almost inevitable when a virus is under selective pressure. We reveal here that P5C3, a previously described potent and broadly active neutralizing mAb that acts by blocking the Spike-ACE2 interaction, not only targets a highly conserved region of the RBD but also can bind Omicron Spike concomitantly with P2G3. We further identify mutants capable of escaping neutralization by either one of these mAbs, but demonstrate that i) they are lowly infectious, ii) they are extremely rare in the wild, suggesting poor fitness, and iii) P5C3 and P2G3 efficiently neutralize each other’s escapees, validating their use in combination. It is additionally noteworthy that P5C3 and P2G3 both efficiently neutralize the now prevalent R346K-containing BA.1.1 and BA.2 Omicron sub-variants, which partly or completely escape blockade by other currently available mAbs.

Many of the spectacular advances gained in the fight against the pandemics, including COVID-19 vaccines and potent neutralizing mAbs, were largely erased in a matter of months following the emergence of the Delta and Omicron variants. The Omicron variant exhibits markedly reduced sensitivity to vaccine-induced humoral immunity in healthy donors but, more importantly, immunocompromised individual are now almost completely unprotected due to their inability to mount a protective humoral immune response following vaccination ^33^. For these vulnerable individuals, we propose that passive immunization through two-to-three injections per year with the extended half-life P2G3 and P5C3 LS mAbs ^27^, that can simultaneously bind to highly conserved and distinct epitopes on the viral Spike, represents a very attractive prophylaxis option^34^. With potent neutralization, Fc-mediated functional activity and demonstrated *in vivo* protection, this broadly active combination, subject to successful development and authorization, has the potential to be a superior anti-SARS-CoV-2 mAb cocktail for prophylactic and therapeutic interventions against all current VOCs, and its breadth of activity suggests that it might be capable of neutralizing many future SARS-CoV-2 VOCs.

## Extended data

**Extended data Fig 1.**
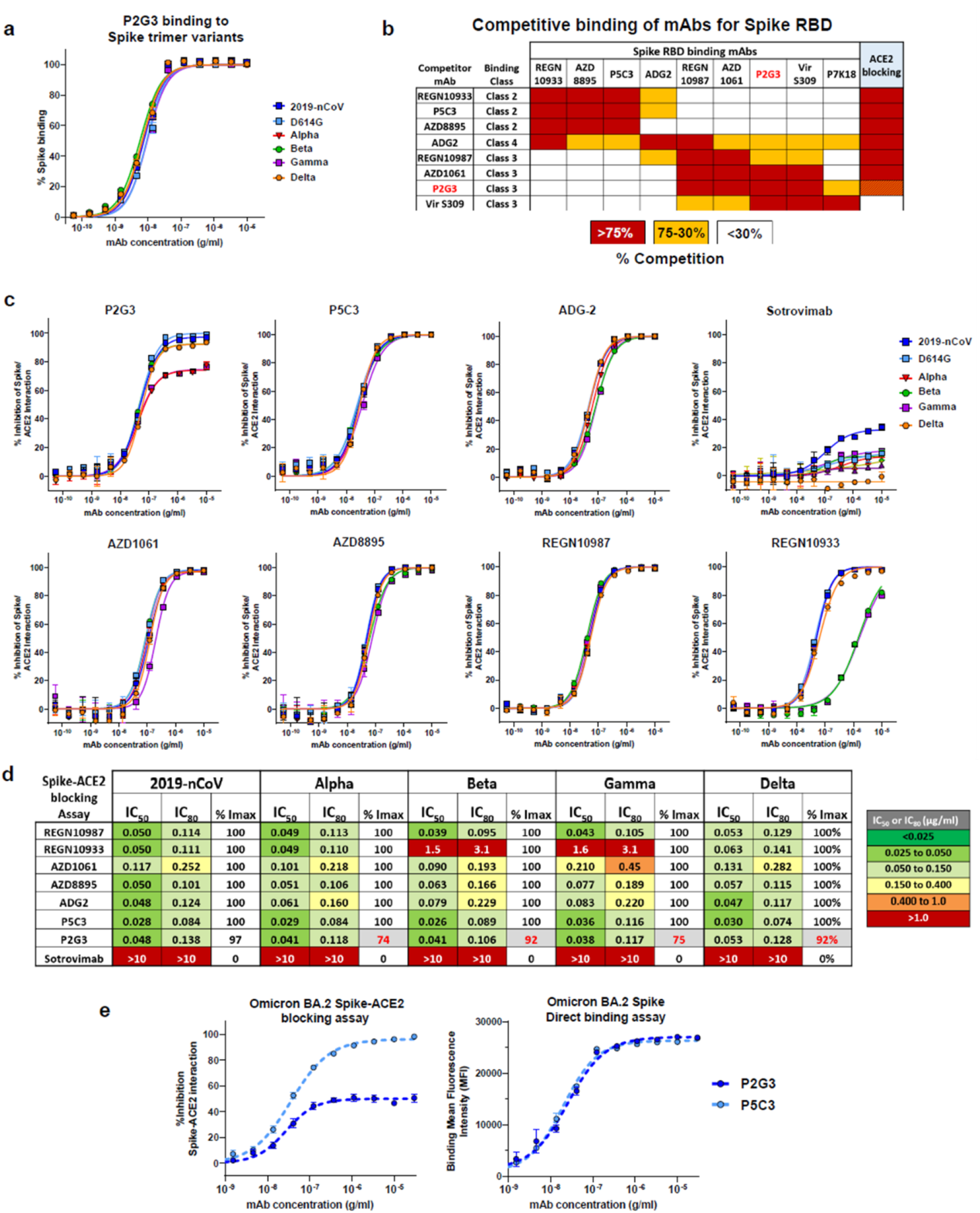
Binding properties of P2G3 and other anti-SARS-CoV-2 antibodies for recombinant Spike trimer proteins from 2019-nCoV and variants of concern in direct binding and Spike-ACE2 interaction assays. **a)** Spike binding curves and **b**) Competitive binding studies between antibodies binding to the 2019-nCoV Spike RBD protein. RBD coupled beads pre-incubated with saturating concentrations of competitor antibody were used for binding studies with mAbs or ACE2. Competitors induced either strong blocking (Red boxes), partial competition (orange boxes) or non-competitive (white boxes) binding with the corresponding mAb to RBD. Red and yellow-hashed lines indicate incomplete blocking of the Spike-ACE2 interaction with Alpha, Gamma and Omicron Spike variant proteins. **c**) Spike-ACE2 blocking activity of a panel of in-house, authorized and clinically advanced anti-Spike mAbs and **d**) heatmap showing IC_50_, IC_80_ and Imax values for our panel of mAbs in the Spike-ACE2 assay. These Luminex based assays were performed with beads coupled with Spike trimer proteins from the original 2019-nCoV, D614G mutant, Alpha, Beta, Gamma, and Omicron SARS-CoV-2 variants of concern. Sotrovimab was included as a control mAb that binds the RBD without blocking the Spike-ACE2 interaction. **e**) Representative data for Spike-ACE2 blocking activity and Spike binding for P2G3 and P5C3 mAbs using the same Omicron BA.2 Spike trimer coated Luminex beads. Data presented is representative of 2-4 independent experiments with each concentration response tested in duplicate. Mean values ± SEM are shown.

**Extended data Fig 2:**
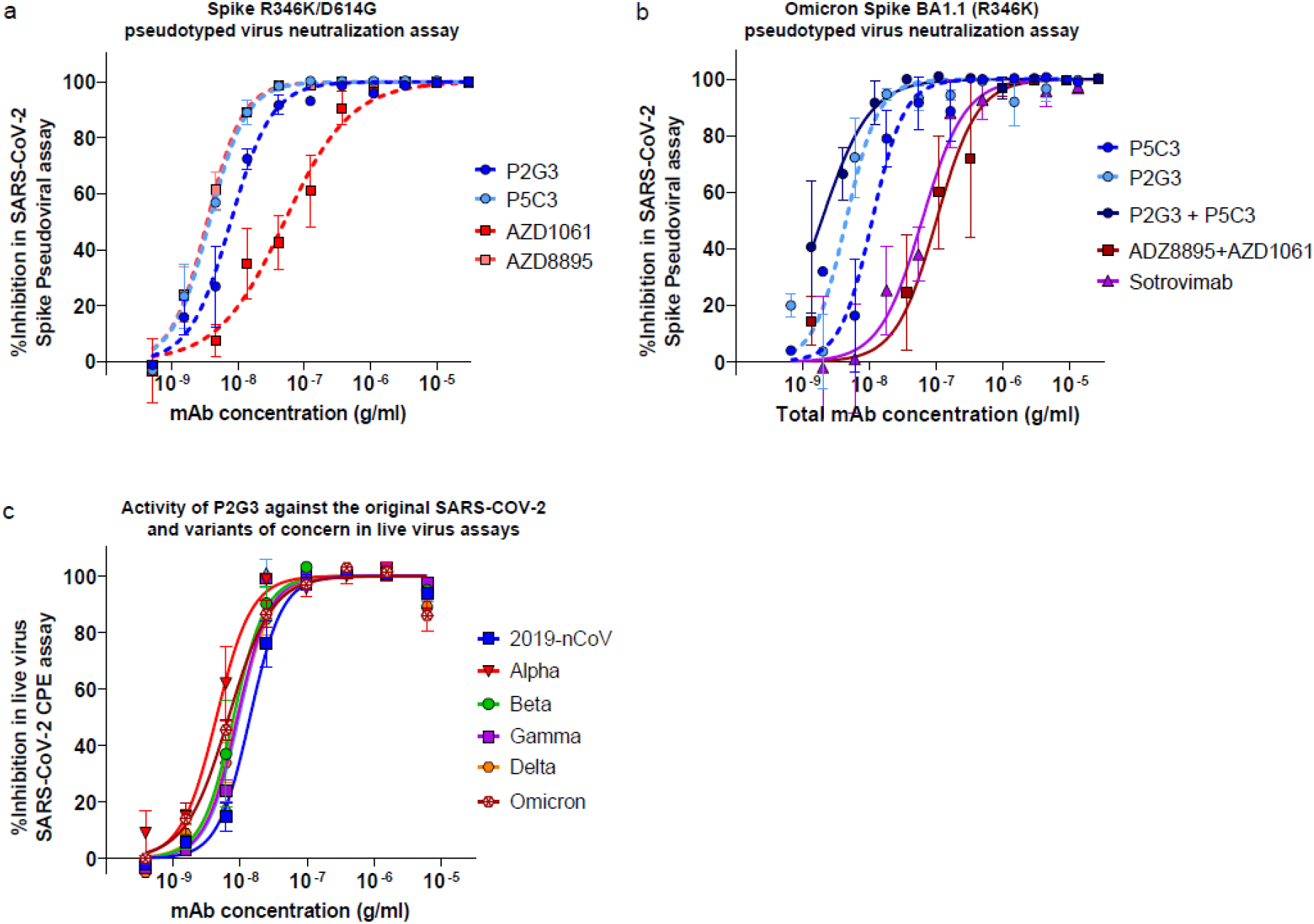
P2G3 retains neutralizing activity against R346K Spike-coated pseudoviruses and all SARS-CoV-2 variants in a live virus cytopathic effect assay. Neutralization of lentiviral particles pseudotyped with: **a)** SARS-CoV-2 Spike encoding R346K mutation and the D614G substitution that became dominant early in the pandemic and **b)** Omicron BA1.1 with the R346K substitution in a 293T-ACE2 infection assay. **c)** P2G3 evaluated in live virus cytopathic effect neutralization assays with the original 2019-nCoV and Alpha, Beta, Gamma, Delta and Omicron BA.1 variants of concern. Results shown are representative of two independent experiments in pseudovirus assays with each concentration response tested in triplicate. Live virus results are representative of 2-4 experiments with each concentration tested in duplicate. Mean values ± SEM are shown.

**Extended data Figure 3.**
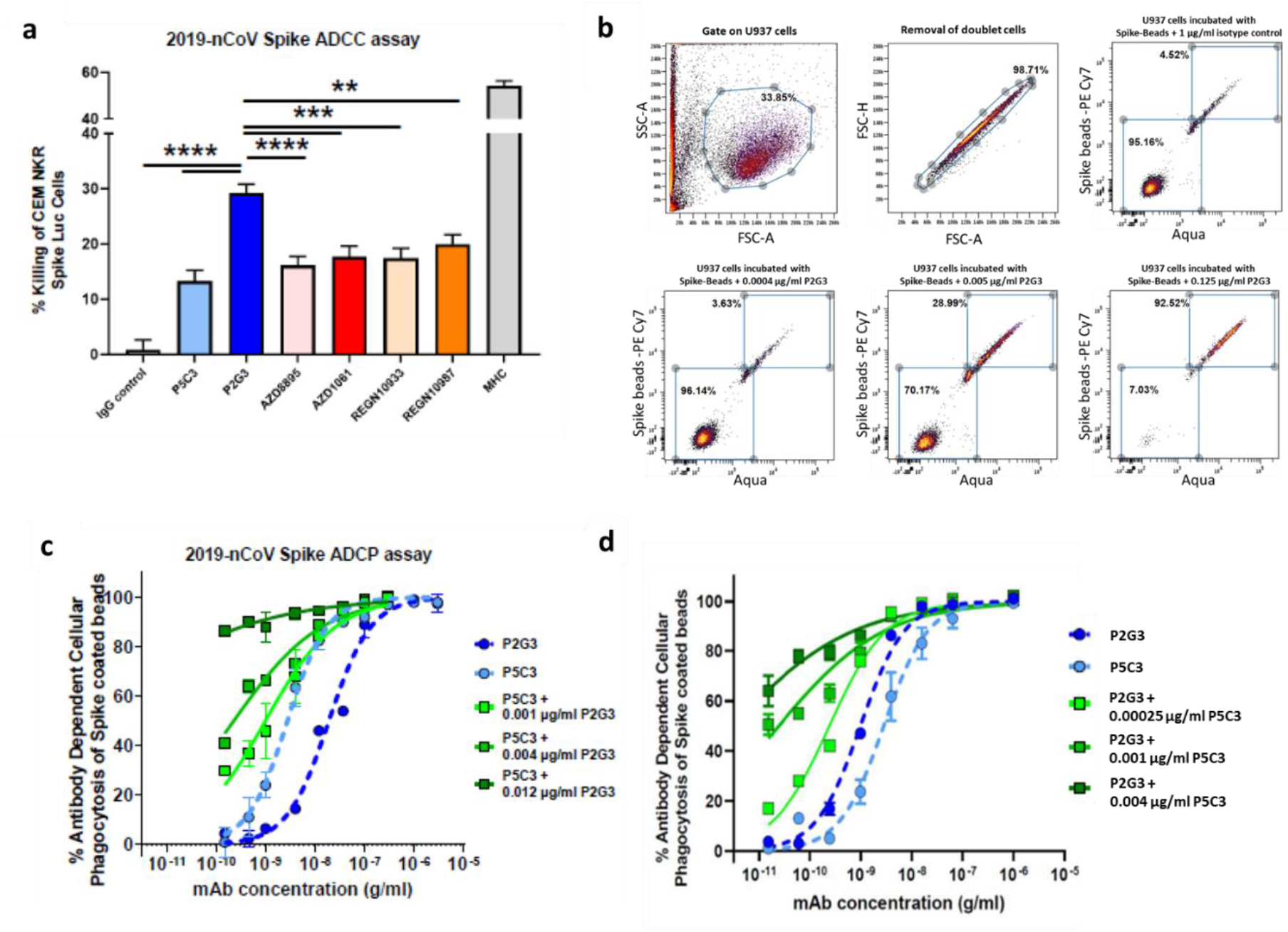
P2G3 LS shows strong Fc-mediated functional activity in ADCC cell killing and ADCP assays. **a)** Antibody dependent cellular cytotoxicity assay (ADCC) performed with CEM NKR Luciferase cells stably expressing cell surface 2019-nCoV Spike. P2G3 exhibits potent ADCC activity in killing Spike positive cells. ADCC experiments performed with five replicates per condition using effector cells from five different healthy donors. CEM NKR Spike cells were incubated with 0.30 μg/ml of the indicated human IgG mAbs. Statistical difference evaluated by Two-way ANOVA with p-values presented as *p<0.05, **p<0.01, ***p<0.001, ****p<0.0001. **b)** Flow cytometry gating strategy for the selection of U937 cells, the removal of cell doublets and the evaluation of cells for which Spike-coated fluorescent beads have undergone phagocytosis. The threshold gate for Spike-specific phagocytosis of beads was set using and isotope control antibody and representative dot plots for P2G3 mediated ADCP of 2019-nCoV Spike-coated beads by U937 are shown with >4000 cells analysed per condition. **c-d)** Antibody dependent cellular cytotoxicity assay performed ancestral 2019-nCoV and with Omicron BA.1 variant Spike protein biotinylated and bound to streptavidin coated fluorescent beads. Beads mixed with the indicated antibody concentrations were incubated with the U937 monocyte effector cell line and antibody dependent cellular phagocytosis of the Spike coated beads was evaluated by flow cytometry. Dashed lines correspond to individual antibodies and solid lines indicate combinations of P2G3 and P5C3. Results shown are representative data for three separate experiments with each concentration response tested in duplicates or triplicates. Mean values ± SEM are shown.

**Extended data Figure 4.**
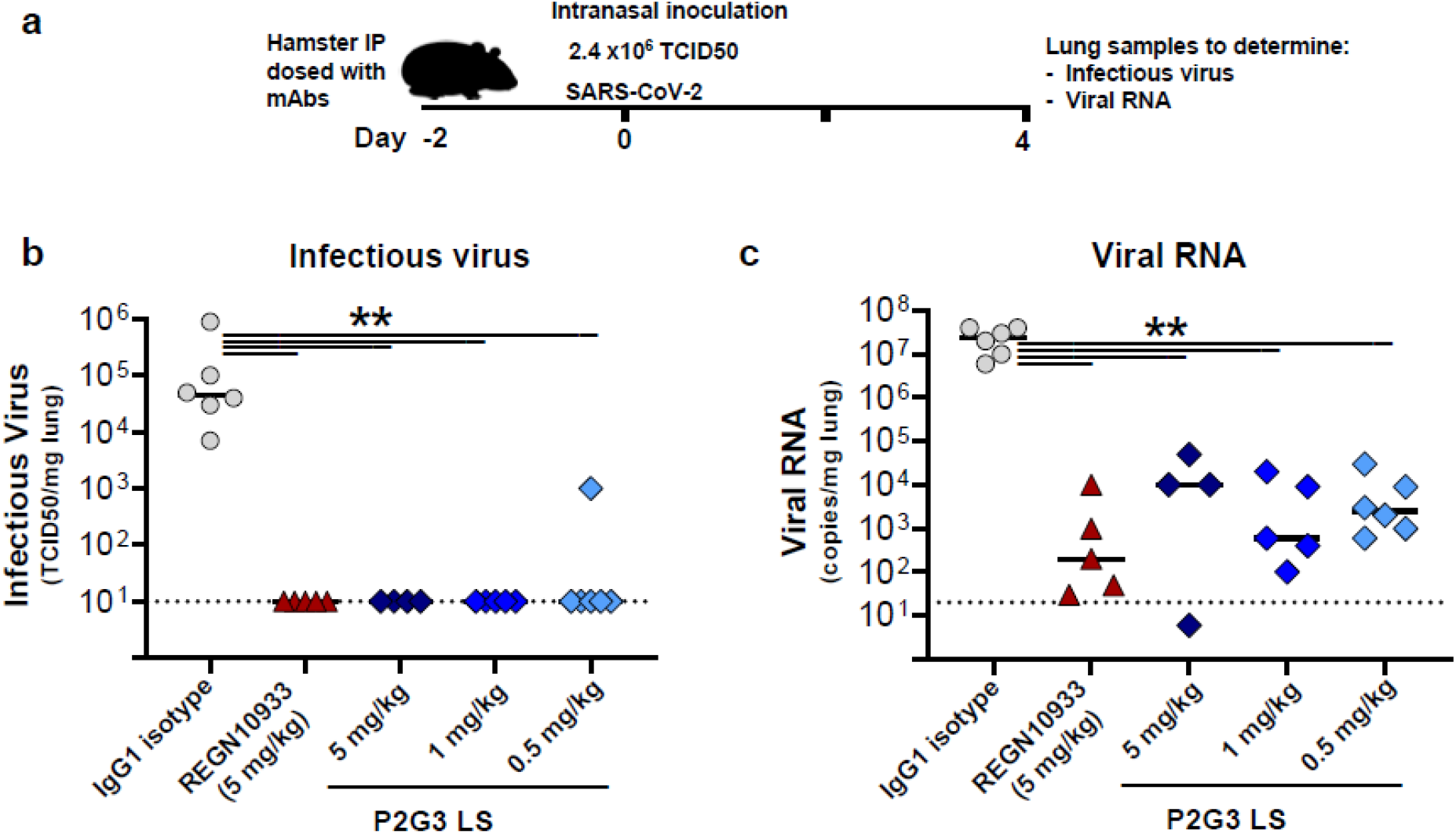
P2G3 LS confers potent *in vivo* efficacy in the hamster challenge model for SARS-CoV-2 infection. **a)** Overview of study design for the SARS-CoV-2 hamster challenge model. Animals were administered intraperitoneally 5.0, 1.0 or 0.5 mg/kg of P2G3, 5 mg/kg of REGN10933 positive control or 5 mg/kg of an IgG1 isotype control and challenged two days later (Day 0) with an intranasal inoculation of the original 2019-nCoV SARS-CoV-2 virus (2.4 ×10^6^ TCID50). **b)** Median levels of infectious virus and **c)** viral RNA copies/mg lung tissue in each of the study arms are shown on day 4 post-inoculation with SARS-CoV-2 virus. A total of 4-6 hamsters were used per P2G3 treatment arm. Non-parametric Mann-Whitney U-tests were used to evaluate the statistical difference between the treatment conditions with p<0.009 (**).

**Extended data Fig 5.**
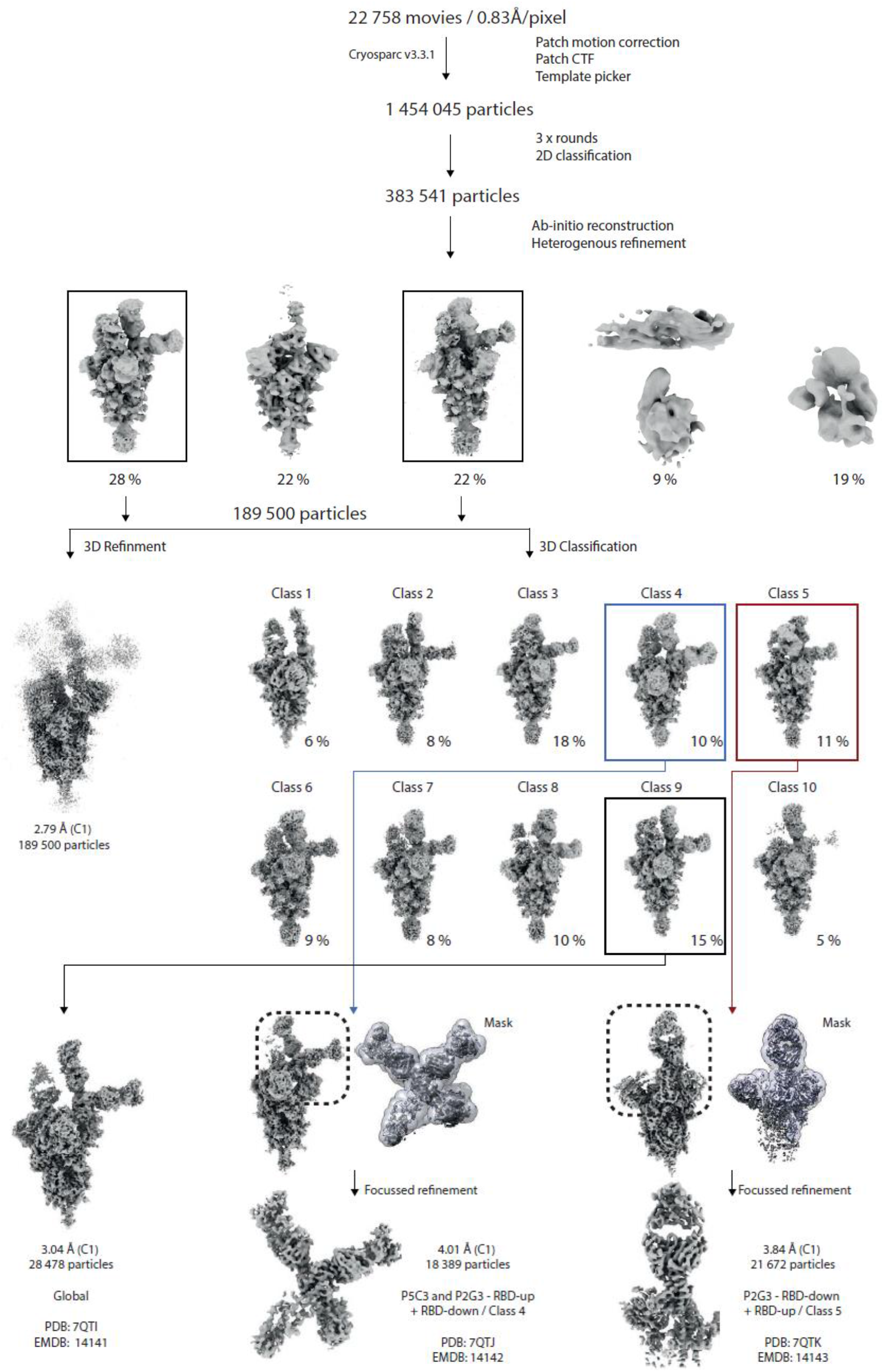
Cryo-EM processing of the Omicron Spike. Cryo-EM processing workflow performed in CryoSPARC v.3.3.1.

**Extended data Fig 6.**
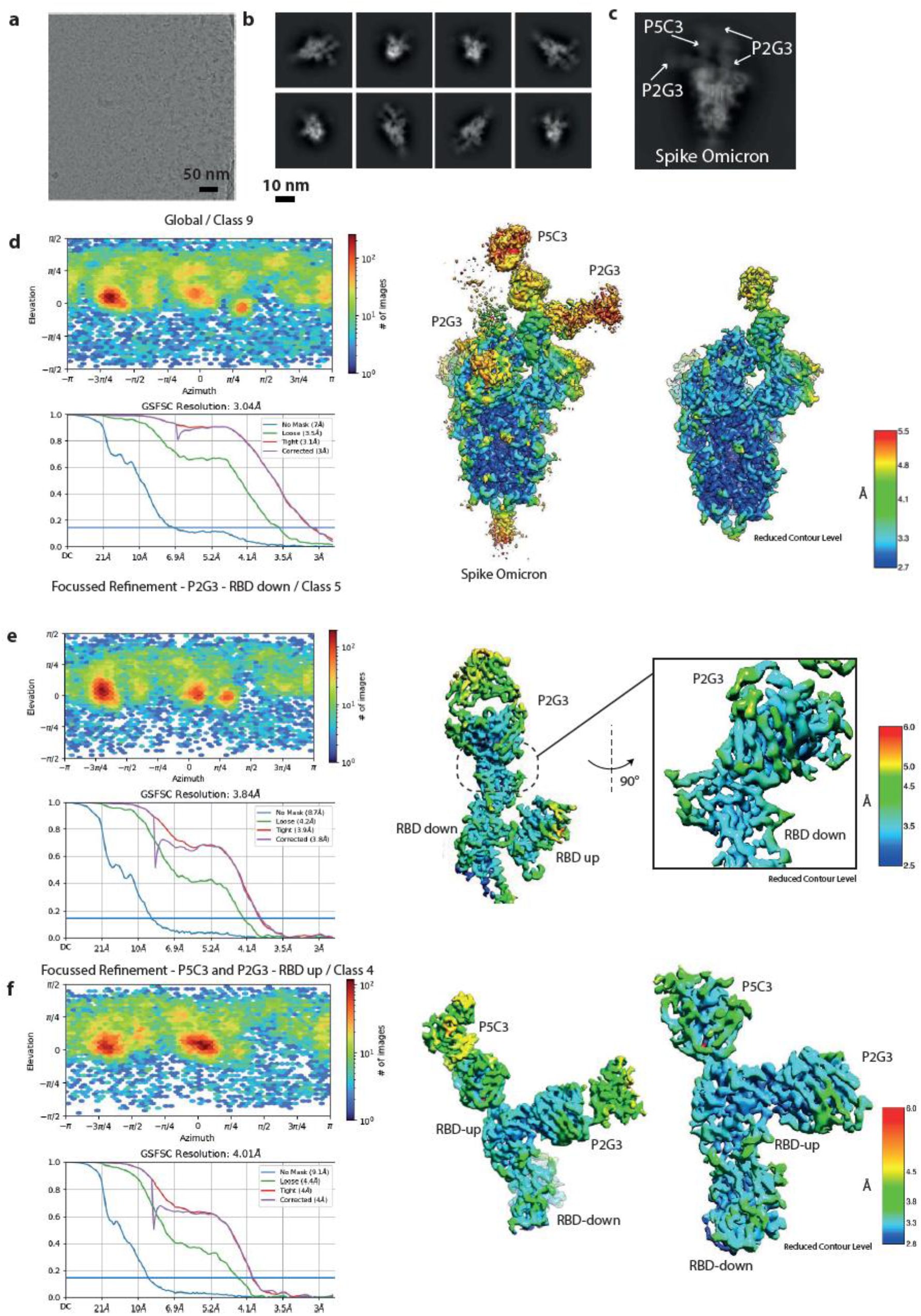
Details of Cryo-EM processing and Resolution of Cryo-EM maps. **a)** Raw representative micrograph. **b)** Representative 2D class averages. **c)** Enlarged 2D class showing the Omicron Spike with bound Fabs. **d)** Direction distribution plot and FSC curves indicating a resolution of 3.04 Å (FSC 0.143) of the Global (Class 9) map off the full-length Omicron Spike bound to Fabs. **e)** Direction distribution plot and FSC curves indicating a resolution of 3.84 Å (FSC 0.143) of the P2G3-bound-RBD-down (Class 5) locally refined map **f)** Direction distribution plot and FSC curves indicating a resolution of 4.01 Å (FSC 0.143) of the P5C3-P2G3-bound-RBD-up (Class 4) locally refined map. Global and focused refined maps coloured by local resolution.

**Extended data Fig 7.**
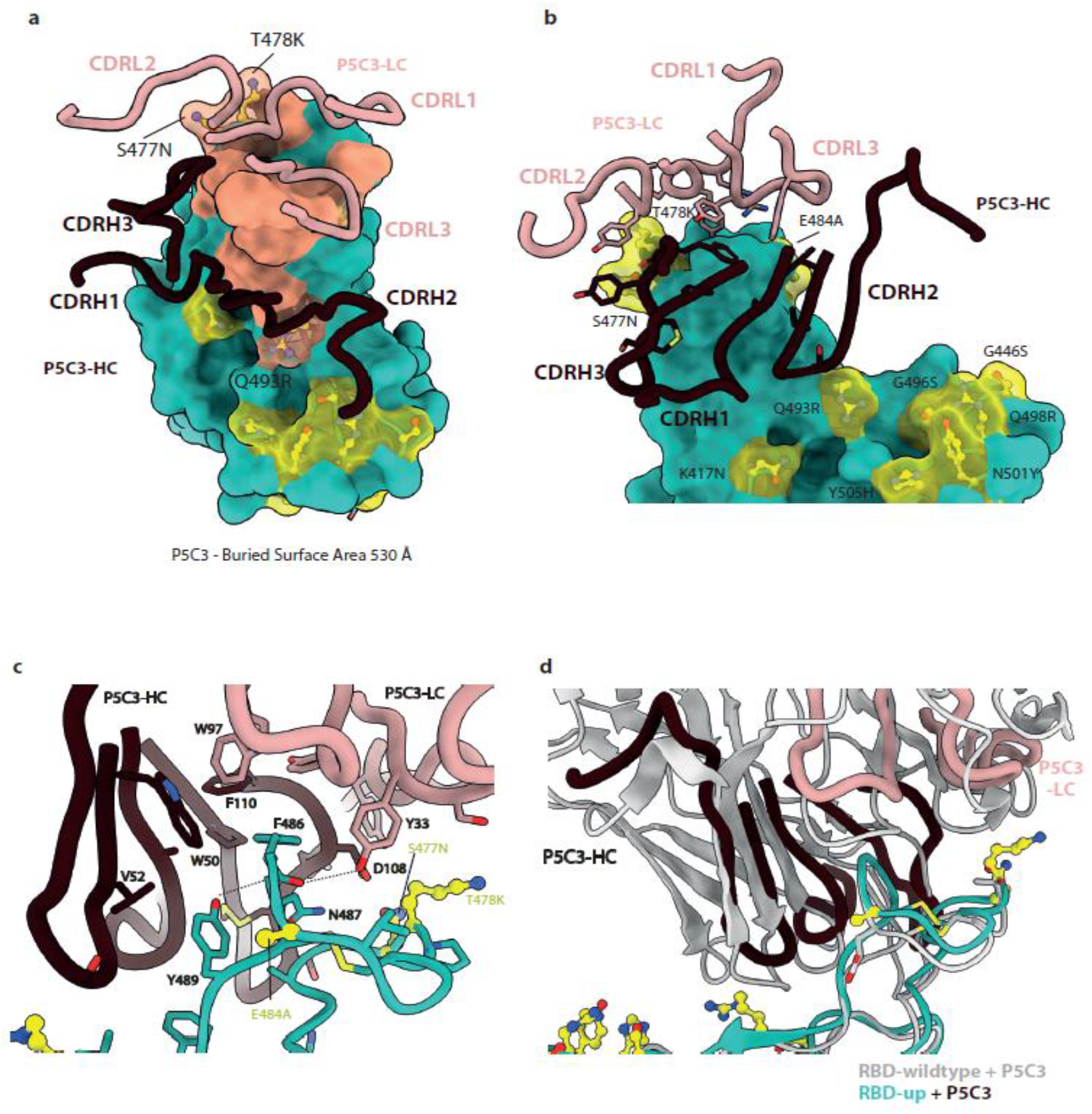
Analysis of the P5C3 Fab-Omicron RBD interacting surfaces. **a)** The buried surface area of P5C3 (pink) overlaid on the RBD surface (green). P5C3 buries around 500 Å of the surface of the Omicron RBD. Specific CDR loops of the heavy and light chains are indicated. Omicron mutations are shown as balls-and-sticks and transparent surfaces in yellow. **b)** Zoomed-in view of the interacting region of P2G3. Specific CDR loops of the heavy and light chains are indicated. Omicron mutations in the region of the Fab are highlighted in yellow. Interacting residues of the Fab are shown as sticks. **c)** Detailed atom level analysis of the interactions between the Omicron RBD shown as ribbons (green) and the P5C3 Fab heavy and light chains shown as liquorice (dark red and pink). Residues at the interface are shown as sticks with potential interactions of interest as dashed lines. Omicron mutations are shown in yellow. **d)** Superposition of the P5C3-Omicron-RBD interface with the P5C3-wild-type-RBD interface (PDB; 7PHG)

**Extended data Fig 8.**
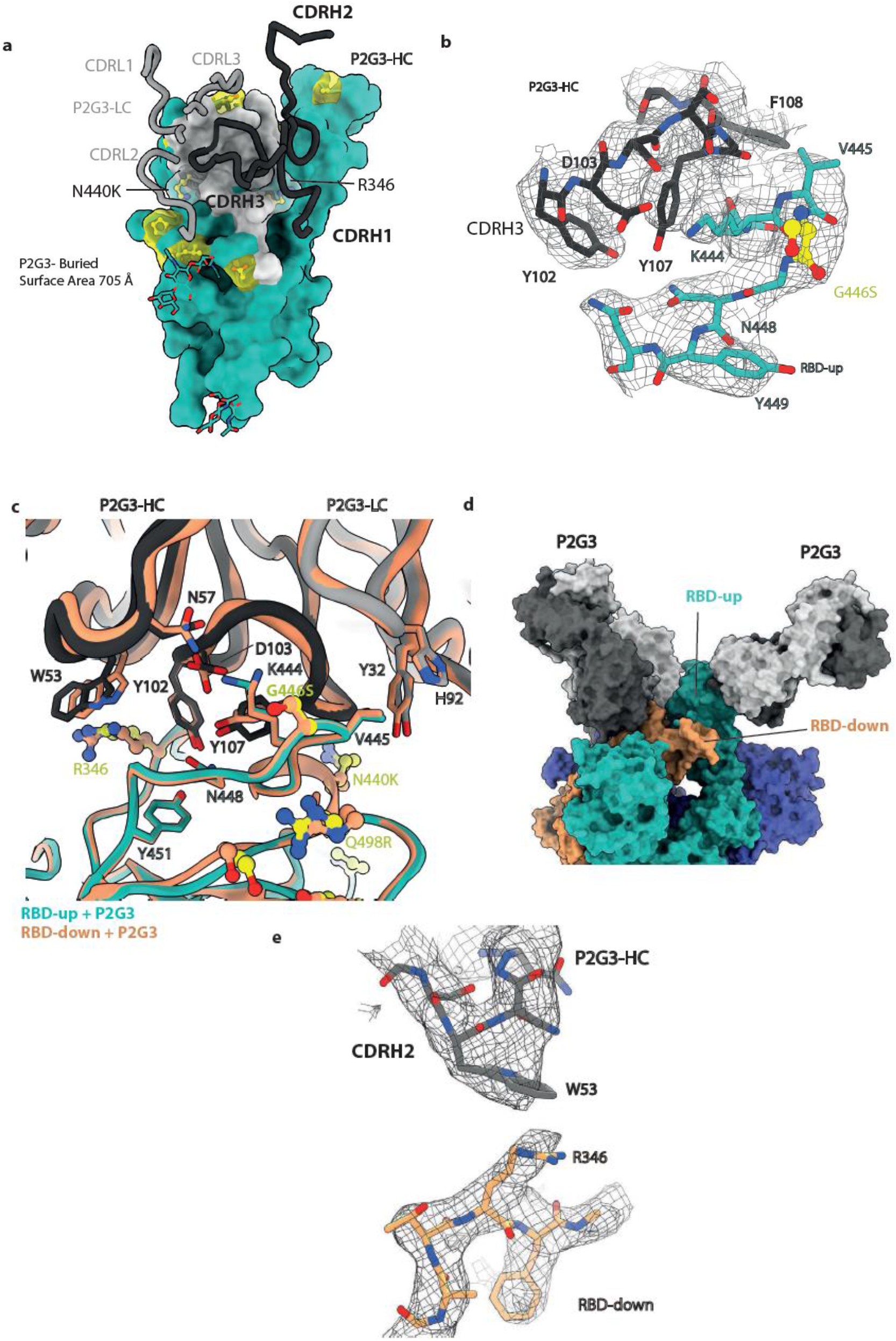
Additional views of the P2G3 Fab-Omicron RBD interacting surfaces. The buried surface area of P2G3 (grey) overlaid on the RBD surface (green). P2G3 buries 705 Å of surface of the Omicron RBD. Specific CDR loops of the heavy and light chains are indicated. Omicron mutations are shown as balls-and-sticks and transparent surfaces in yellow. Stick representation of the P2G3 interface of CDRH3 and the RBD region containing residues 440-451. The mesh represents the Cryo-EM density. **c**) Superposition of the P2G3-RBD-up interface with the P2G3-RBD-down interface shows no significant differences with interactions conserved. **d**) Binding of P2G3 Fabs to the RBD-up domain in green and RBD-down domain in orange within the trimeric Spike. **e**) Stick representation of the P2G3 interface of CDRH2 residue W53 that forms a potential cation-pi interaction with RBD residue R346. The mesh represents the Cryo-EM density.

**Extended data Fig. 9.**
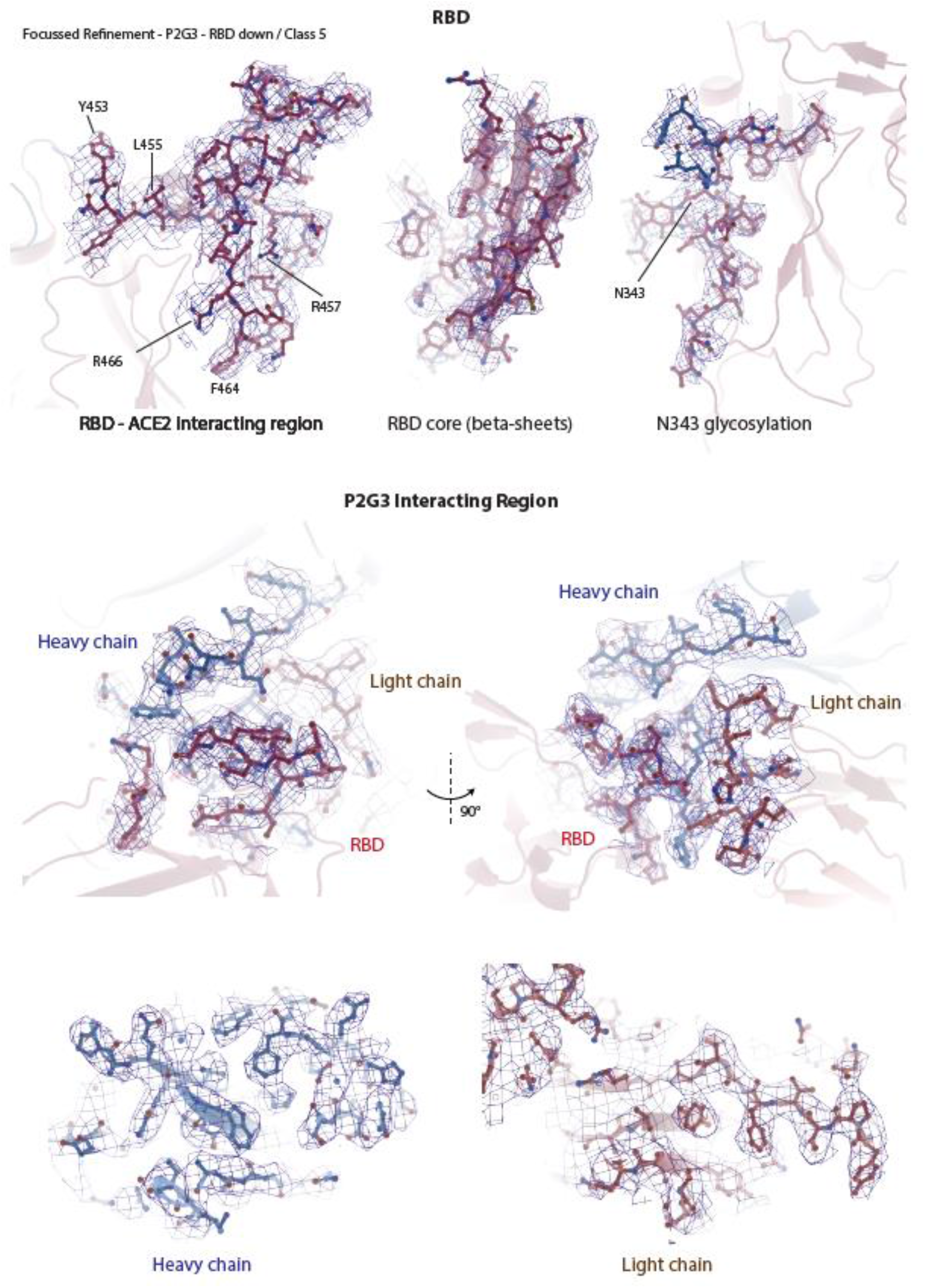
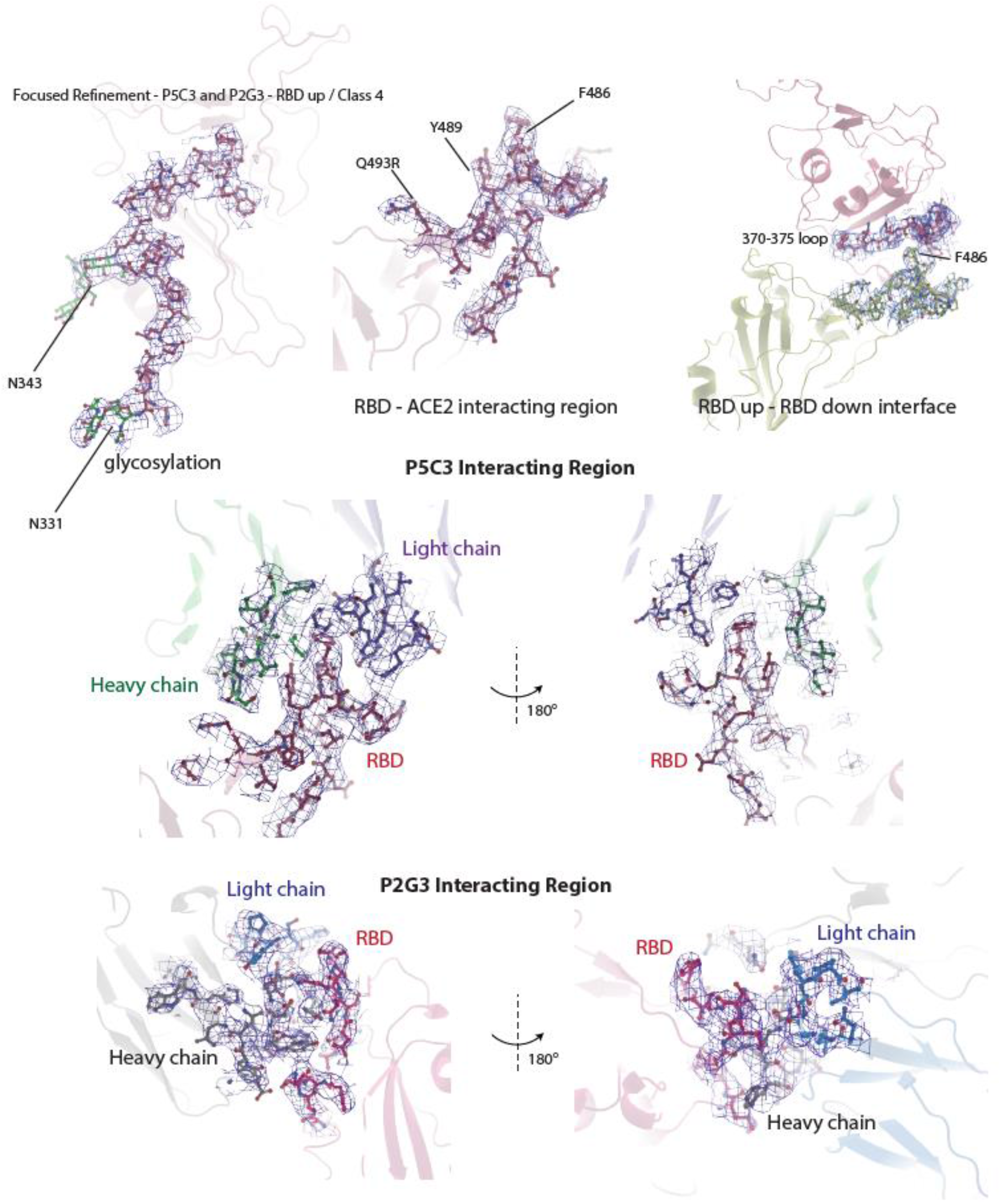
Highlights of regions of the Omicron Spike and Fabs with Cryo-EM density maps. The Cryo-EM density is rendered as a mesh. The atomic model is shown as ribbon or stick representation.

**Extended data Figure 10.**
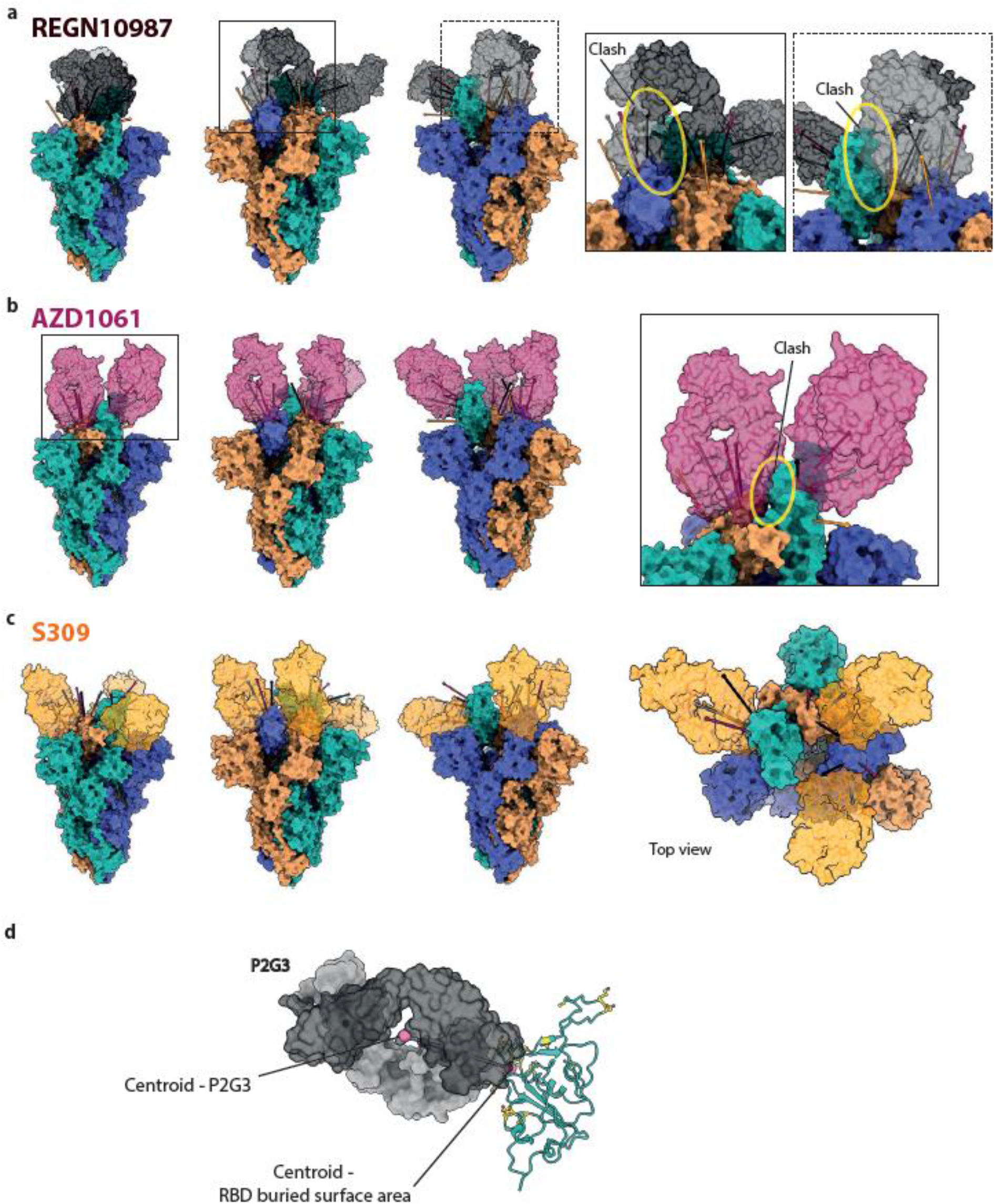
Angle of attack of Fabs for authorized and clinically advanced anti-SARS-CoV-2 mAbs modelled to the Omicron Spike trimer. Model of Fabs for Class 3 antibodies, REGN10987, AZD1061 and S309/Sotrovimab binding each of the RBD protomers for the full Omicron trimer. Trimers are shown from multiple perspectives to visualize the different angles of attack the Fabs have depending on the Spike protomer. **a)** REGN10987 Fab bind to the green RBD-up conformation but modelled REGN10987 Fab bound to either the RBD-down of the orange or blue protomer would clash sterically with the adjacent blue RBD-down or green RBD-up protomers, respectively. Together it is predicted REGN10987 is able to bind only RBD-up. **b)** AZD1061 is predicted to bind the RBD-up form and RBD-down form of the blue protomer but AZD1061 Fab modelled binding to the RBD-down of the orange protomer would potentially clash with the adjacent green RBD-up. **c)** S309/Sotrovimab is able to bind all RBDs simultaneously as shown for P2G3. Angle of attack of Fabs to the RBD is defined as the line connecting the centroid of the Fab to the centroid of the surface area of the RBD that the Fabs bury.

**Extended data Figure 11.**
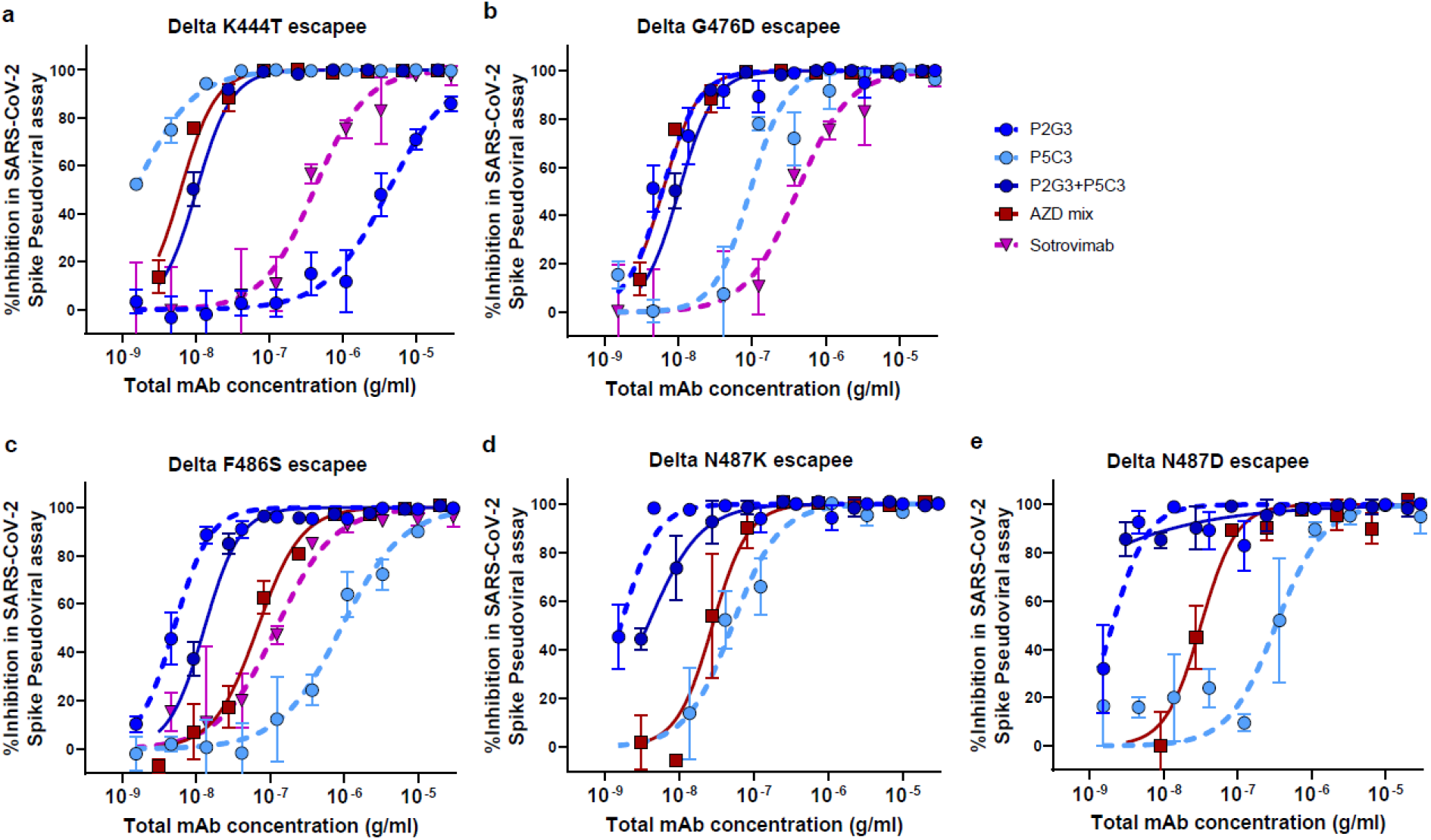
P2G3 and P5C3 efficiently cross-neutralize Spike-coated pseudoviruses with mutations encoding one another’s escapees. Neutralization of lentiviral particles pseudotyped with Delta variant SARS-CoV-2 Spike encoding: a) the P2G3-escaping K444T Spike substitution and P5C3-escaping b) G476D c) F486S, d) N487K and N487D Spike substitutions. Results shown are representative of one experiment in pseudovirus assays with each concentration response tested in triplicate. Mean values ± SEM are shown.

**Extended data Table 1.** Cryo-EM data collection, refinement and validation statistics.

|  | SARS-CoV-2 S<br>Omicron Spike<br>B.1.1.529-P2G3-P5C3<br>(Global map) | SARS-CoV-2 S Omicron Spike<br>B.1.1.529-P2G3 bound to RBD-down + RBD-up<br>(Local map) | SARS-CoV-2 S Omicron Spike<br>B.1.1.529-P2G3-P5C3 bound to<br>RBD-up + RBD-down<br>(Local map) |
| --- | --- | --- | --- |
| **Microscope** | TFS Titan Krios G4 + E-CFEG |  |  |
| **Detector** | Falcon 4 |  |  |
| Magnification | 165kx |  |  |
| Voltage (kv) | 300 |  |  |
| Electron exposure (e-/Å<sup>2</sup>) | 60 |  |  |
| Defocus range (um) | -0.8 to -2.5 |  |  |
| Pixel size (Å) | 0.83 |  |  |
| Symmetry | C1 |  |  |
| Micrographs | 22 758 |  |  |
| Initial particle images (No.)<br>(After manual 2D class<br>curation) | 383 540 |  |  |
| Final particle images (No.) | 28 478 | 21 672 | 18 839 |
| Map resolution (Å) (*FSC*<br>*0.143*) | 3.04 | 3.84 | 4.01 |
| Map resolution range (Å) | 25.9-2.76 | 10.5-3.02 | 30.0-3.02 |
| **Refinement** |  |  |  |
| Model resolution (Å) (*FSC*<br>*0.5*) | 3.72 | 4.14 | 4.38 |
| Initial model used | 7QO7 | --- | --- |
| Map sharpening B factor (Å<sup>2</sup>) | -24.4 | -38.5 | -44.1 |
| Protein residues | 5 061 | 643 | 1 280 |
| **RMSD deviations** |  |  |  |
| Bond lengths (Å) [#Z>5] | 0.003 [0] | 0.002 [0] | 0.003 [1] |
| Bond angles (°) [#Z>5] | 0.620 [16] | 0.609 [2] | 0.579 [4] |
| **Validation** |  |  |  |
| **MolProbity score** | **1.82** | **1.85** | **1.77** |
| Clashscore | 7.95 | 9.04 | 8.08 |
| % Poor rotamers (%) | 0.02 | 0.00 | 0.00 |
| C-beta outliers (%) | 0.00 | 0.00 | 0.00 |
| **Ramachandran plot** |  |  |  |
| Favored (%) | 94.34 | 94.63 | 95.25 |
| Allowed (%) | 5.46 | 5.21 | 4.51 |
| Disallowed (%) | 0.20 | 0.16 | 0.24 |
| **PDB** | 7QTI | 7QTK | 7QTJ |
| **EMDB** | 14141 | 14143 | 14142 |

## METHODS

### Method details

#### Study COVID-19 donors

Serum and blood mononuclear cell samples were from donors participating in the ImmunoCov and ImmunoVax studies performed by the Immunology and Allergy Service, Lausanne University Hospital. Study design and use of subject samples were approved by the Institutional Review Board of the Lausanne University Hospital and the ‘Commission d’éthique du Canton de Vaud’ (CER-VD).

### Production of SARS-CoV-2 Spike proteins

SARS-Cov2 Spike mutations are similar for all the cloned Spike variants and the corresponding viral isolates and are listed in Table 2. Production of 2019-nCoV (D614G), B.1.17, B.1.351 and P.1 variants has already been described ^21^. RNA isolated from an anonymized leftover sample of an individual suspected to be SARS-CoV-2 Omicron strain infected was reverse transcribed into cDNA. The Omicron Spike ectodomain was amplified by PCR with primers (listed in Table 3) designed on consensus sequence from available Omicron sequences, and introduced by In-Fusion cloning into the nCoV-2P-F3CH2S plasmid, replacing the original wild-type Spike^29^. The 2 prolines (P986-P987) and the furin cleavage site mutations (residues 682-685 mutated to GSAS) stabilizing the Spike protein in the trimeric prefusion state were further introduced simultaneously by PCR and In-Fusion-mediated site directed mutagenesis using primers listed in Table 3 as previously described^2^, and the full Omicron ORF was sequence verified. The Delta B1.617.2 variant clone was generated by gene synthesis with a codon optimized Spike ORF (GenScript). The final constructs encode the Spike ectodomains, containing a native signal peptide, the 2P and furin cleavage site mutations, a C-terminal T4 foldon fusion domain to stabilize the trimer complex followed by C-terminal 8x His and 2x Strep tags for affinity purification. The trimeric Spike variants were produced and purified as previously described^16^. The purity of Omicron Spike trimers used for cryo-EM was determined to be >99% pure by SDS-PAGE analysis. Biotinylation of Spike or RBD proteins was performed using the EZ-Link™ NHS-PEG4-Biotin (Life Technologies) using a 3-fold molar excess of reagent and using the manufacturer’s protocol. Biotinylated proteins were buffer exchanged with PBS using an Amicon Ultra-0.5 with a 3 kDa molecular weight cut-off. Spike and RBD tetramers were prepared fresh before use and formed by combining biotinylated proteins with PE-conjugated Streptavidin (BD Biosciences) at a molar ratio of 4:1.

**Table 2:** Spike mutations.

|  |  |
| --- | --- |
| **Alpha**<br>**B.1.1.7** | Δ69-70, Δ144, N501Y, A570D, D614G, P681H, T716I, S982A, D1118H |
| **Beta**<br>**B.1.351** | L18F, D80A, D215G, Δ242-244, R246I, K417N, E484K, N501Y, D614G, A701V |
| **Gamma**<br>**P.1** | L18F, T20N, P26S, D138Y, R190S, K417T, E484K, N501Y, D614G, H655Y, T1027I, V1176F |
| **Delta**<br>**B.1.617.2** | T19R, Δ156-157, R158G, L452R, T478K, D614G, P681R, D950N |
| **Omicron**<br>**B.1.1.529.1** | A67V, Δ69-70, T95I, G142D, Δ143-145, Δ211, L212I, ins214EPE, G339D, S371L, S373P, S375F, K417N, N440K, G446S, S477N, T478K, E484A, Q493R, G496S, Q498R, N501Y, Y505H, T547K, D614G, H655Y, N679K, P681H, N764K, D796Y, N856K, Q954H, N969K, L981F |

**Table 3:**
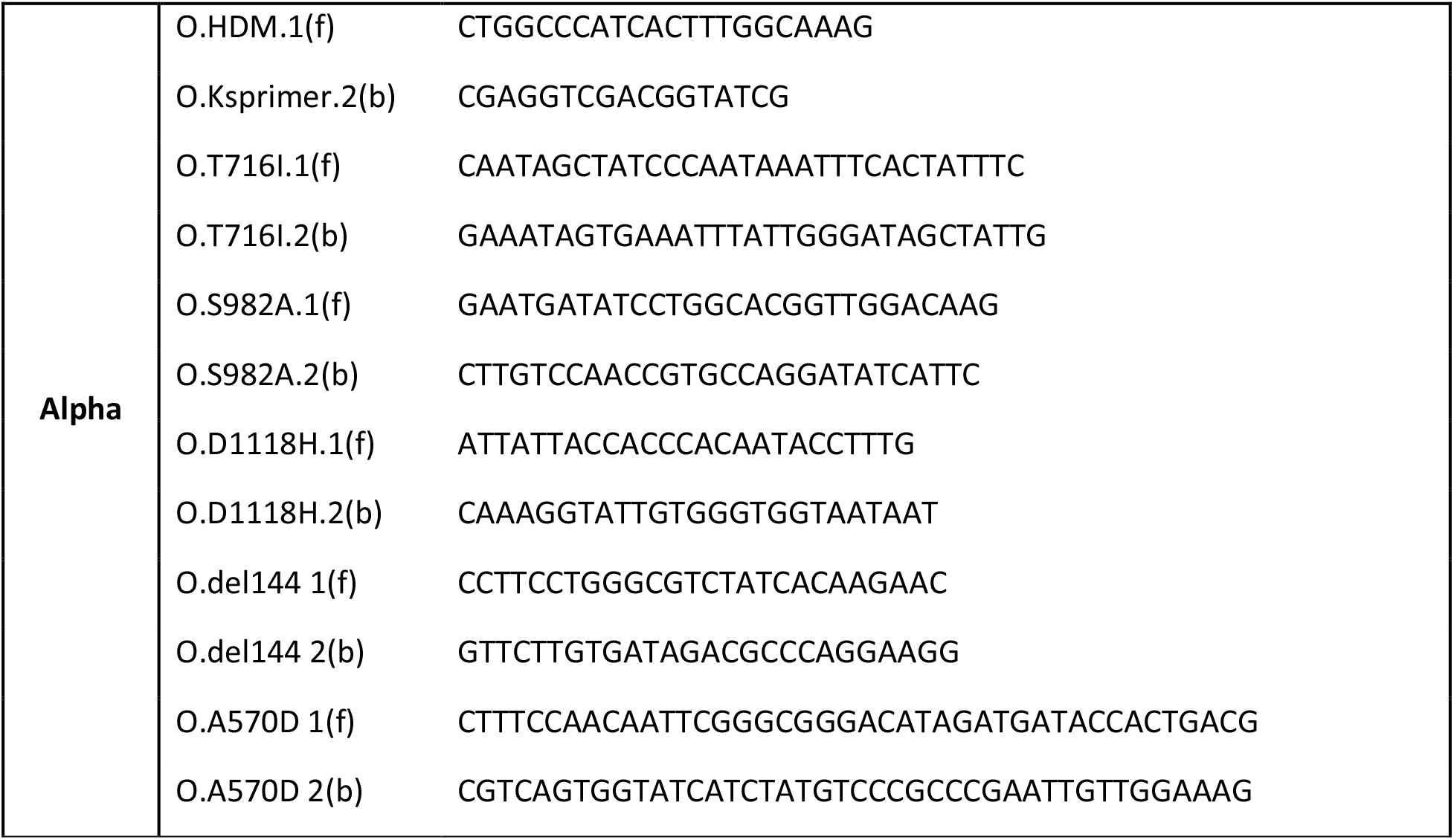
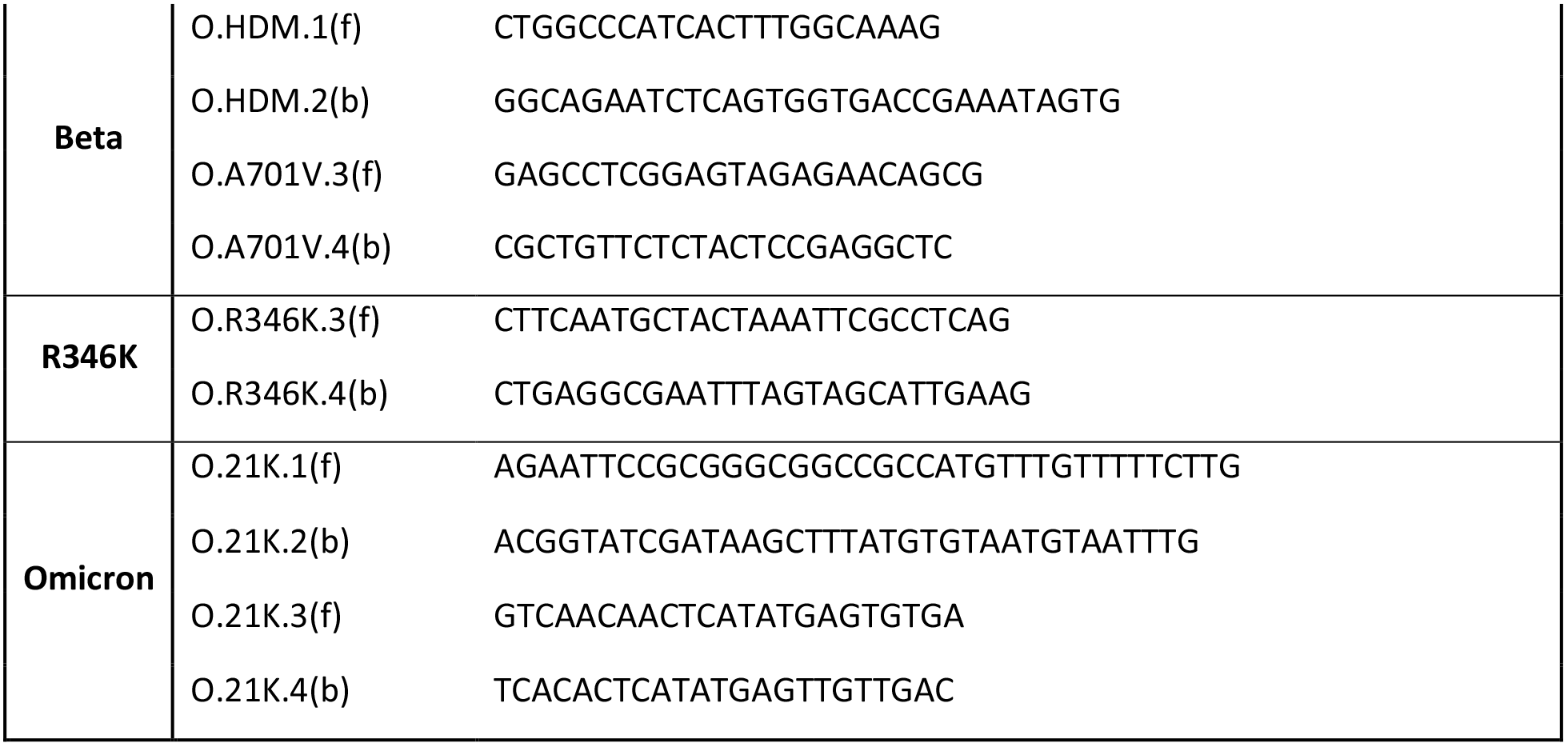
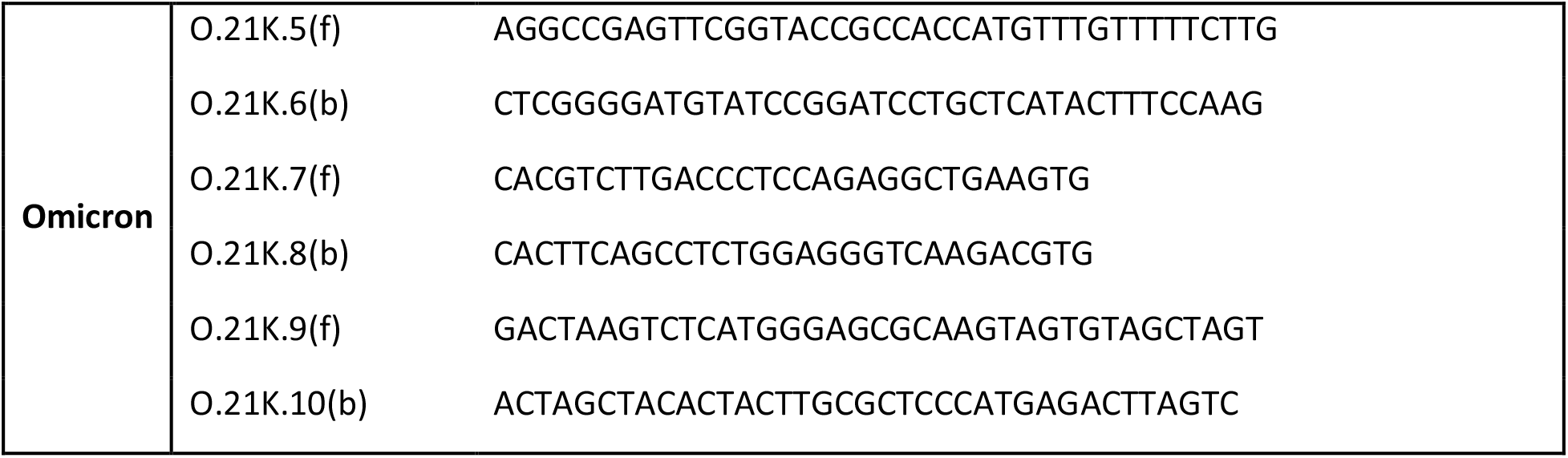
Primers used to clone spike mutations.

### Binding and ACE2 blocking studies with SARS-CoV-2 Spike

Luminex beads used for the serological and purified antibody binding assays were prepared by covalent coupling of SARS-CoV-2 proteins with MagPlex beads using the manufacturer’s protocol with a Bio-Plex Amine Coupling Kit (Bio-Rad, France). Each of the SARS-CoV-2 Spike proteins expressed with different mutations were coupled with different colored MagPlex beads so that tests could be performed with a single protein bead per well or in a multiplexed Luminex binding assay. Binding curves for antibody affinity measurements and the Spike-ACE2 interaction assay were performed as previously described ^16,35^. Competitive binding studies were performed by pre-incubating 25 μg/ml of the indicated competitor antibody with the original 2019-nCoV RBD protein coupled Luminex beads for 30 minutes. Biotinylated P5C3, P2G3, REGN10933, REGN10987, AZD8895, AZD1061, ADG-2 or S309 antibodies (prepared as described above) were added to each well at 1 μg/ml followed by a further 20-minute incubation. Biotinylated antibody bound to RBD in the presence of competitor was stained with Streptavidin-PE at a 1:1000 dilution (BD Biosciences) and analysed on a 200 Bioplex instruments. COVID-19 serum samples from >100 donors were monitored for levels of IgG antibody binding to the SARS-CoV-2 Spike trimer proteins from 2019-nCoV, D614G, Alpha, Beta and Gamma in the Luminex bead based assay.

### Anti-Spike B cell sorting, immortalization and cloning

The blood from a ImmunoVax study donors were collected in EDTA tubes and the isolation of blood mononuclear cell was performed using Leucosep centrifuge tubes (Greiner Bio-one) prefilled with density gradient medium (Ficoll-PaqueTM PLUS, GE Healthcare) according to the manufacturer’s instructions. Freshly isolated cells were stained with the cocktail of fluorescent conjugated antibodies containing anti-CD19 APC-Cy7, anti-CD3-BV510, anti-IgM-FITC, anti-IgD PE-CF594, anti-CD27-APC, anti-CD38-V450 (BD Biosciences) along with the pre-complexed Beta variant Spike tetramer (2 μg in 100μl) coupled to PE-streptavidin (BD Biosciences). All other aspects with cell sorting, immortalization and cloning were as described in Fenwick et al ^21^.

### SARS-CoV-2 live virus stocks

All the biosafety level 3 procedures were approved by the Swiss Federal Office of Public Health. The SARS-Cov2 D614G isolate and B.1.1.7 clone have previously been described ^21^. Beta (EPI_ISL_981782), Gamma (EPI_ISL_981707), Delta B1.617.2 (EPI_ISL_1811202) and Omicron B.1.1.529.1 (EPI_ISL_7605546) early isolates were a kind gift from I. Eckerle, Geneva University Hospitals. Viral stocks were prepared in EPISERF on VeroE6 or Calu-3 (for Omicron), aliquoted, frozen and titrated on VeroE6 cells.

### SARS-CoV-2 live virus cell based cytopathic effect neutralization assay

Neutralization assay was performed as previously described except for Delta and Omicron isolates were EPISERF instead of DMEM 2% FCS was used to prepare antibodies serial dilutions. Equal amount of different viruses was used in all experiments (1200 plaque forming units per well), except for the less cytopathic Omicron strain, where 2.5 times more virus was incubated with each antibody tested in parallel.

### Selection of resistant virus in presence of mAbs

The day before infection, 293T + ACE2 (+/− TMPRSS2) cells were seeded in 6-well plates coated with poly-Lysin at a density of 1.×10^6^ cells per well. To generate a viral population under mAb pressure, early passage virus was diluted in 1 ml EPISERF 2% FCS and incubated with 0.25ng/ml mAb for 1hrs at 37°C in duplicates. Each mixture was added to the cells and P1 (passage 1) supernatants were harvested 3 days later and clarified on 0.45um SpinX filters centrifuged at 4000×g for 4 minutes. Aliquots of cleared P1 supernatants were diluted 1:40 in DMEM 2%, incubated with mAbs as described above and used to infect fresh cells for 4 days. P2 supernatants were treated as P1 and P3 supernatants were collected for RNA extraction and subsequent selection step. To select for mAb resistant viruses, 100 μl of the cleared undiluted P3 heterogeneous viral population was incubated with 100 μl mAbs at 2.5 μg/ml, 0.625 μg/ml or 0.155 μg/ml final concentration for 1hrs at 37°C. Mixture was then applied on cells in 400 μl DMEM 2% (1:2 volume) for 3 to 4 days. Viruses were propagated for a few more passages and aliquots of each passage was used for RNA extraction and sequencing. Virus produced in absence of mAb was collected and treated the same way in parallel to control for appearance of mutations due to cell culture conditions.

### Spike-pseudotyped lentivectors production and neutralization assays

HDM-IDTSpike-fixK plasmid (BEI catalogue number NR-52514, obtained from J.D. Bloom, Fred Hutchinson Cancer Research Center) was used as backbone for all the clonings. For Alpha and Beta clones, the HDM-IDTSpike-fixK NotI/SmaI fragments were swapped with the respective Alpha and Beta fragments from the previously described pTwist plasmids^21^. Alpha P681H, T716I, S982A, D1118H and Beta A701V were further added in respective ORFs as well as R346K in D614G plasmid by In-Fusion-mediated site directed mutagenesis using primers described in Table 3. Delta B1.617.2 clone was generated by gene synthesis with a codon-optimized Spike ORF (GenScript). The Omicron ORF was amplified from an RNA as described for protein production with primers listed in Table 3. Pseudoviruses were alternatively produced with the original 2019-nCoV (Cat #100976), Alpha / B.1.1.7 (Cat #101023) and Beta/B.1.351 (Cat #101024) pCAGGS-SARS2-Spike vectors obtained from NIBSC. These vectors were co-transfected with pMDL p.RRE, pRSV.Rev and pUltra-Chili-Luc vectors (Addgene) into 293T cells in DMEM medium + 10% FCS using Fugene 6 (Promega) for pseudoviruses production. Neutralization assays were performed as previously described ^21^.

### NHP challenge model for SARS-COV-2 Omicron BA.1 infection

Cynomolgus macaques (Macaca fascicularis) originating from Mauritian AAALAC certified breeding centers were used in this study. All animals were housed within IDMIT animal facilities at CEA, Fontenay-aux-Roses under BSL-2 and BSL-3 containment when necessary (Animal facility authorization #D92-032-02, Préfecture des Hauts de Seine, France) and in compliance with European Directive 2010/63/EU, the French regulations and the Standards for Human Care and Use of Laboratory Animals, of the Office for Laboratory Animal Welfare (OLAW, assurance number #A5826-01, US). Animals tested negative for Campylobacter, Yersinia, Shigella and Salmonella before being use in the study.

The protocols were approved by the institutional ethical committee “Comité d’Ethique en Expérimentation Animale du Commissariat à l’Energie Atomique et aux Energies Alternatives” (CEtEA #44) under statement number A20-011. The study was authorized by the “Research, Innovation and Education Ministry” under registration number APAFIS#24434-2020030216532863. All information on the ethics committee is available at https://cache.media.enseignementsup-recherche.gouv.fr/file/utilisation_des_animaux_fins_scientifiques/22/1/comiteethiqueea17_juin2013_257221.pdf.

Four female cynomolgus macaques aged 3–6 years were randomly assigned between the control and treated groups to evaluate the efficacy of P2G3 LS in the prophylaxis challenge study. The treated group (n = 2 [MF1 and MF2]) received one dose at 10 mg/kg of P2G3 LS human IgG1 monoclonal antibody delivered by intravenous injection three day prior to challenge, while control animals (n = 2 in parallel [MF3 and MF4] and n=2 historical [MF5 and MF6]) received no treatment. All animals were then exposed to a total dose of 10^5^ TCID50 of Omicron B.1.1.529 SARS-CoV-2 virus produced in Calu-3 cells (NIH/BEI reference: NR-56462) via the combination of intranasal and intratracheal routes (day 0) with sample collection and testing performed as previously described ^36^.

### Hamster challenge model SARS-CoV-2 infection

Hamster studies were performed at KU LEUVEN R&D as previously described ^21^. A pathologist at KU LEUVEN R&D performed lung histology at Day 4 post-challenge with individual scores attributed to: 1) lung congestion, 2) intra alveolar haemorrhage, 3) apoptotic bodies in bronchus wall, 4) necrotizing bronchiolitis, 5) perivascular edema, bronchopneumonia, 6) perivascular inflammation, 7) peribronchial inflammation and 8) vasculitis. Cumulative pathology scores showed that all challenged hamsters in IgG control groups had positive histological signs of pathology, while the P2G3 and REGN10933 treatment groups showed no significantly different pathology scores relative to average cumulative scores for non-infected animals.

### Antibody Fc-mediated functional activity assays

Antibody dependent cellular cytotoxicity (ADCC) and antibody dependent phagocytosis (ADCP) assays were performed as previously described with minor changes ^37^. Cryopreserved peripheral blood mononuclear cells from healthy patient were thawed and resuspended at 1 million/ml in RPMI medium (Gibco, Life Technologies) supplemented with 10% heat-inactivated fetal bovine serum (FBS), 100 IU/ml penicillin and 100 μg/ml streptomycin (BioConcept). Cells were stimulated with 25 ng/ml IL-15 (Miltenyi Biotec) for 6 hours and ADCC effector cells were enriched from PBMCs by depletion of T cells using anti-CD3 coupled magnetic beads from the EasySep™ Human T Cell Isolation Kit (Stemcell). Target cells used for the ADCC assay were a CEM-NK resistant cell line that was stably transfected to express the original 2019-nCoV Spike protein at the cell surface and with constitutive expression of the Luciferase gene (CEM-NKR-Spike-Luc cells). In the ADCC assay, CEM-NKR-Spike-Luc cells were incubations with anti-Spike antibody at 0.3 μg/ml, isotype control antibodies at 0.3 μg/ml or an anti-HLA class I (MHC) positive control antibody (Invivogen) at 0.005 μg/ml. Following 5 minutes at room temperature, the CEM-NKR-Spike-Luc cells / antibody mixes were then co-cultured overnight at a 1:10 ratio of CD3-depleted PBMC effector cells in RPMI medium supplemented with 5% low-IgG FBS and 1% penicillin-streptomycin in U-bottom 96-well plate (SARSTEDT). The following day, cell killing was monitored either directly by flow cytometry or indirectly by monitoring the decrease in luciferase activity associated with cell death. In flow cytometry analysis, Spike-transfected CEM NKR-Luc cells were stained with PKH26 kit according the manufacturers protocol (Sigma) prior to performing the ADCC assay. To monitor cell killing, co-cultured cells were washed and stained with fluorescent conjugated antibodies, CD56-AF488 (BD Biosciences,), CD16-FITC (BD Biosciences), Aqua Live/Dead cell stain (Invitrogen), CD4-PECF594 (BD Biosciences) and Annexin V-APC (BD Biosciences), and then analysed using a FACS LSR II cytometer instrument. Spike CEM-NKR-Luc cells were gated on with the PKH26 fluorochrome and then dead (Aqua positive) and apoptotic/dying (AnnexinV positive) cells were evaluated for the positive (anti-HLA class I), negative (isotope control) and test (anti-Spike antibodies) antibody conditions to establish ADCC activity. In the luciferase readout assay, co-cultured cells were transferred in white-Elmer 96-well plate and luciferase activity were measured using One-Step Luciferase assay kit (BPS Biosciences) on a Synergy plate reader instrument. Co-cultured Spike CEM-NKR-Luc cells incubated with isotope control antibodies generally gave 5-10% reduced luminescence signal compared to CEM-NKR-Luc cells incubated in the absence of effector cells. Positive control anti-MHC (anti-HLA class I) antibody gave strong antibody dependent cell killing of 60-80% and anti-Spike antibodies gave intermediate responses.

In the ADCP assay, TransFluoSpheres™ Carboxylate-Modified Microspheres (1.0 μm diameter, 488nm/560nm excitation/emission, ThermoFisher) were coupled directly with 2019-nCoV trimeric Spike protein or with streptavidin according to the manufactures protocol. Spike coupled beads were washed, incubated in the presence or absence of the different concentrations of anti-Spike antibody for 30 minutes and then the mix was added directly to the U937 monocyte cell line plated in a 96-well U-bottom 96-well plate (Sarstedt). Following an overnight incubation, cells were analysed using a FACS LSR II cytometer instrument to identify cells with Spike bead fluorescence. U937 cells incubated with Spike beads in the absence of antibody generally showed <5% phagocytic activity while increased ADCP activity was observed with increasing concentration of anti-Spike antibody with a maximum of 100% of cells exhibiting fluorescence associated with Spike bead phagocytosis. ADCP activity of Omicron Spike coated beads was performed by pre-incubating Streptavidin coupled TransFluoSpheres™ beads with biotinylated Omicron Spike protein, produced as described above. Beads were washed after 30 minutes and used in the ADCP assay as described for the directly coupled 2019-nCoV trimeric Spike beads.

### Cryo-electron microscopy

Cryo-EM grids were prepared with a Vitrobot Mark IV (Thermofisher Scientific (TFS)). Quantifoil R1.2/1.3 Au 400 holey carbon grids were glow-discharged for 120 s at 15mA using a PELCO easiGlow device (Ted Pella, Inc.). 3.0 μL of a 0.7 mg/ml Omicron Spike mixed with 0.16 mg/mL each of P5C3 and P2G3 Fab fragments (Final 3.2 uM Omicron Spike:1.5 uM P5C3:1.5 uM P2G3) was applied to the glow-discharged grids, and blotted for 6 s under blot force 10 at 100% humidity and 4 °C in the sample chamber, and the blotted grid was plunge-frozen in liquid nitrogen-cooled liquid ethane.

Grids were screened for particle presence and ice quality on a TFS Glacios microscope (200kV), and the best grids were transferred to TFS Titan Krios G4. Cryo-EM data was collected using TFS Titan Krios G4 transmission electron microscope (TEM), equipped with a Cold-FEG on a Falcon IV detector in electron counting mode. Falcon IV gain references were collected just before data collection. Data was collected using TFS EPU v2.12.1 using aberration-free image shift protocol (AFIS), recording 4 micrographs per ice hole.

Movies were recorded at magnification of 165kx, corresponding to the 0.83A pixel size at the specimen level, with defocus values ranging from −0.8 to −2.5 μm. Exposures were adjusted automatically to 60 e^−^/Å^2^ total dose, resulting in an exposure time of approximately 3 seconds per movie. In total, 22 758 micrographs in EER format were collected.

### Cryo-EM image processing

On-the-fly processing was first performed during data acquisition for evaluating the data quality during screening by using cryoSPARC live v3.3.1 ^38^. The obtained ab-initio structures were used for better particle picking for template creation. Motion correction was performed on raw stacks without binning using the cryoSPARC implementation of motion correction^39^. 1 454 045 particles were template-picked automatically picked. Three rounds of 2D classification were performed, resulting in a particle set of 383 541 particles. Selected particles resulting from the 2D classification were used for ab-initio reconstruction and hetero-refinement. After hetero-refinement, 189 500 particles contributed to an initial 3D reconstruction of 2.79 Å resolution (FSC 0.143) with C1 symmetry. These particles were subjected to 3D classification resulting in 10 classes. Class 9 resulted in a global map of the Omicron Spike with an RBD-up bound to a P5C3 and P2G3 Fab at a resolution of 3.04 Å (FSC 0.143) with C1 symmetry. Focussed refinement of Class 4 with a soft mask volume encompassing an RBD-up and its bound Fab and an adjacent RBD-down resulted in a map at 4.01 Å (FSC 0.143) with C1 symmetry. Finally, focussed refinement of Class 5 with a soft mask volume encompassing an RBD-down its bound P2G3 and an adjacent NTD resulted in a map at 3.84 Å (FSC 0.143) with C1 symmetry. The soft mask volume were generated manually in UCSF Chimera and cryoSPARC ^40^.

### Cryo-electron microscopy model building

A model of a Spike trimer (PDB ID 7QO7) or AlphaFold2 (ColabFold implementation) models of the P5C3 and P2G3 Fabs were fit into the cryo-EM maps with UCSF Chimera These docked models were extended and rebuilt manually with refinement, using Coot and Phenix^41,42^. Figures were prepared in UCSF Chimera, UCSF ChimeraX and Pymol^40^. Numbering of the full-length Spike models within the global map is based on Omicron numbering. Numbering of models containing just the RBD within the local maps are based on wild-type numbering. Fab numbering of both heavy and light chains start at one from the CH1 and CL domains respectively. Buried surface area measurements and centroid measurements were calculated within ChimeraX.

### Statistical analysis

Statistical parameters including the exact value of n, the definition of center, dispersion, and precision measures (Mean or Median ± SEM) and statistical significance are reported in the Figures and Figure Legends. Data were judged to be statistically significant when p < 0.05. In Figures, asterisks denote statistical significance as calculated using the two-tailed non-parametric Mann-Whitney U test for two groups’ comparison. Analyses were performed in GraphPad Prism (GraphPad Software, Inc.) and Microsoft Excel.

## DATA AVAILABILITY

The reconstructed maps of the global Omicron Spike with Fabs bound are available from the EMDB database, C1 symmetry, EMDB-14141. The atomic model for the full-length Omicron Spike with Fabs bound is available from the PDB database, PDB-7QTI. The local focussed-refinement map of the RBD-up with two Fabs bound is available from the EMDB database, EMDB-14142. The atomic model for the RBD-up with two Fabs bound in the locally refined map is available from the PDB database, PDB-7QTJ. The local focussed-refinement map of the RBD-down with P2G3 Fab bound is available from the EMDB database, EMDB-14143. The atomic model for the RBD-down with P2G3 Fab bound in the locally refined map is available from the PDB database, PDB-7QTK. All plasmids made in this study are available upon request to the corresponding authors.

## Acknowledgements

We thank the Service of Immunology and Allergy at the Lausanne University Hospital for analysis of serum samples for levels of anti-Spike protein IgG antibodies. We thank Isabella Eckerle, Meriem Bekliz and the Virology laboratory of Geneva University Hospital for the Omicron RNA sample and variant isolates collection, the Geneva Genome Center for sequencing and Julien Duc for the development of in house scripts for analyses. We thank Laurence Durrer, Rosa Schier, Michaël François and Soraya Quinche from the EPFL Protein Production and Structure Core Facility for mammalian cell production and purification of proteins, Alexander Myasnikov, Bertrand Beckert and Sergey Nazarov from the Dubochet Center for Imaging (an EPFL, UNIGE, UNIL initiative) for cryoEM grids preparation and data collection and Davide Demurtas for set up of cryo-EM condition for automated acquisition in related studies not included in this manuscript. We would also like to thank David Wyatt and members of the CARE-IMI work package 4 team for helpful discussions. G.P. received a grant from the Corona Accelerated R&D in Europe (CARE) project funded by the Innovative Medicines Initiative 2 Joint Undertaking (JU) under grant agreement No 101005077. The JU receives support from the European Union’s Horizon 2020 research and innovation program, the European Federation of Pharmaceutical Industries Associations (EFPIA), the Bill & Melinda Gates Foundation, Global Health Drug Discovery Institute and the University of Dundee. The content of this publication only reflects the author’s view and the JU is not responsible for any use that may be made of the information it contains. Additional funding was provided through the Lausanne University Hospital (to G.P.), the Swiss Vaccine Research Institute (to G.P. and NCCR TransCure to H.S.), Swiss National Science Foundation Grants (to G.P.) and through the EPFL COVID fund (to D.T.).

## Author contributions

C.F. designed the strategy for isolating and profiling anti-Spike antibodies, designed the functional assays, coordinated the research activities, analysed the data, wrote the initial draft and contributed to the editing of the manuscript. P.T. established designed and performed the experiments with live SARS-CoV-2 virus cytopathic effect neutralization assay, designed engineered and tested the Spike protein mutations and cloningwith the help of C.R., analysed the results and contributed to the editing of the manuscript. L.P., D.N., K.L., F.P. and H.S. coordinated the cryo-EM analysis, analysed the structural data, wrote the manuscript structural section and contributed to the editing of the manuscript. Other contributed as follows: L.E.-L., performed the B cells sorting, immortalization, binding studies and mAb functional assays; A.F. and E.L., cloning of cloned mAb VH and HL; J.C., binding studies, production of select lentiviruses and pseudoviral assays; C.P., in vitro characterization of serum antibodies from donor samples; F.F. mAb purification, mAb characterization and molecular biology.; C.R. performed site directed mutagenesis of the Spike constructs; F.P. coordinated production of recombinant Spike protein and mAb. P.L., Y.L. and R.L. designed the in vivo study, which was executed by C.H, R.M., R.A., C.S.F., G.V. and J.N. G.P. and D.T. conceived the study design, analysed the results and wrote the manuscript.

## Competing Interest Statement

C.F., G.P., P.T. and D.T. are co-inventors on a patent application that encompasses the antibodies and data described in this manuscript (EP 22153464.7 and PCT/IB2022/050731). DT and GP are amongst the founders of and own equity in Aerium Therapeutics, which has rights to and is pursuing the development of the antibodies described in the publication, and has Sponsored Research Agreements with the Lausanne University Hospital (CHUV) and the Ecole Polytechnique Fédérale de Lausanne (EPFL).

## References

1. WHO. Weekly epidemiological update on COVID-19 - 18 January 2022. Vol. July 12th, 2020 (ed. 75, E.) (2022).

2. Elbe, S. & Buckland-Merrett, G. Data, disease and diplomacy: GISAID’s innovative contribution to global health. Glob Chall 1, 33–46 (2017).

3. Viana, R., et al. Rapid epidemic expansion of the SARS-CoV-2 Omicron variant in southern Africa. Nature (2022).

4. Barnes, C.O., et al. Structures of Human Antibodies Bound to SARS-CoV-2 Spike Reveal Common Epitopes and Recurrent Features of Antibodies. Cell 182, 828–842 e816 (2020).

5. Piccoli, L., et al. Mapping Neutralizing and Immunodominant Sites on the SARS-CoV-2 Spike Receptor-Binding Domain by Structure-Guided High-Resolution Serology. Cell 183, 1024–1042 e1021 (2020).

6. Wang, P., et al. Antibody resistance of SARS-CoV-2 variants B.1.351 and B.1.1.7. Nature 593, 130–135 (2021).

7. Garcia-Beltran, W.F., et al. Multiple SARS-CoV-2 variants escape neutralization by vaccine-induced humoral immunity. Cell 184, 2372–2383 e2379 (2021).

8. Iketani, S., et al. Antibody evasion properties of SARS-CoV-2 Omicron sublineages. Nature (2022).

9. Cao, Y.W., J.; Jian, F.; Xiao, T.; Song, W.; Yisimayi, A.; Huang, W.; Li, Q.; Wang, P.; An, R.; Wang, J.; Wang, Y.; Niu, X.; Yang, S.; Liang, H.; Sun, H.; Li, T.; Yu, Y.; Cui, Q.; Liu, S.; Yang, X.; Du, S.; Zhang, Z.; Hao, X.; Shao, F.; Jin, R.; Wang, X.; Xiao, J.; Wang, Y.; Xie, X.S.;. Omicron escapes the majority of existing SARS-CoV-2 neutralizing antibodies. Biorxiv (2022).

10. VanBlargan, L.A., et al. An infectious SARS-CoV-2 B.1.1.529 Omicron virus escapes neutralization by therapeutic monoclonal antibodies. Nat Med (2022).

11. Liu, L., et al. Striking Antibody Evasion Manifested by the Omicron Variant of SARS-CoV-2. Nature (2021).

12. Mannar, D., et al. SARS-CoV-2 Omicron variant: Antibody evasion and cryo-EM structure of spike protein-ACE2 complex. Science, eabn7760 (2022).

13. Planas, D., et al. Considerable escape of SARS-CoV-2 Omicron to antibody neutralization. Nature (2021).

14. Wang, Z., et al. mRNA vaccine-elicited antibodies to SARS-CoV-2 and circulating variants. Nature 592, 616–622 (2021).

15. Chen, R.E., et al. Resistance of SARS-CoV-2 variants to neutralization by monoclonal and serum-derived polyclonal antibodies. Nat Med (2021).

16. Fenwick, C., et al. A high-throughput cell- and virus-free assay shows reduced neutralization of SARS-CoV-2 variants by COVID-19 convalescent plasma. Sci Transl Med 13 (2021).

17. Baum, A., et al. Antibody cocktail to SARS-CoV-2 spike protein prevents rapid mutational escape seen with individual antibodies. Science 369, 1014–1018 (2020).

18. Dong, J., et al. Genetic and structural basis for SARS-CoV-2 variant neutralization by a two-antibody cocktail. Nat Microbiol 6, 1233–1244 (2021).

19. Rappazzo, C.G., et al. An Engineered Antibody with Broad Protective Efficacy in Murine Models of SARS and COVID-19. bioRxiv (2020).

20. Pinto, D., et al. Cross-neutralization of SARS-CoV-2 by a human monoclonal SARS-CoV antibody. Nature 583, 290–295 (2020).

21. Fenwick, C., et al. A highly potent antibody effective against SARS-CoV-2 variants of concern. Cell Rep 37, 109814 (2021).

22. Lempp, F.A., et al. Lectins enhance SARS-CoV-2 infection and influence neutralizing antibodies. Nature 598, 342–347 (2021).

23. Yamin, R., et al. Fc-engineered antibody therapeutics with improved anti-SARS-CoV-2 efficacy. Nature 599, 465–470 (2021).

24. Schafer, A., et al. Antibody potency, effector function, and combinations in protection and therapy for SARS-CoV-2 infection in vivo. J Exp Med 218 (2021).

25. Winkler, E.S., et al. Human neutralizing antibodies against SARS-CoV-2 require intact Fc effector functions for optimal therapeutic protection. Cell 184, 1804–1820 e1816 (2021).

26. Yu, Y., et al. Antibody-dependent cellular cytotoxicity response to SARS-CoV-2 in COVID-19 patients. Signal Transduct Target Ther 6, 346 (2021).

27. Zalevsky, J., et al. Enhanced antibody half-life improves in vivo activity. Nat Biotechnol 28, 157–159 (2010).

28. Ni, D.L., K.; Turelli, P.; Raclot, C.; Beckert, B.; Nazarov, S.; Pojer, F.; Myasnikov, A.; Stahlberg, H.; Trono, D.;. Structural analysis of the Spike of the Omicron SARS-COV-2 variant by cryo-EM and implications for immune evasion. Biorxiv (2021).

29. Wrapp, D., et al. Cryo-EM structure of the 2019-nCoV spike in the prefusion conformation. Science 367, 1260–1263 (2020).

30. Sztain, T., et al. A glycan gate controls opening of the SARS-CoV-2 spike protein. Nat Chem 13, 963–968 (2021).

31. Halfmann, P.J., et al. SARS-CoV-2 Omicron virus causes attenuated disease in mice and hamsters. Nature (2022).

32. Abdelnabi, R., et al. The omicron (B.1.1.529) SARS-CoV-2 variant of concern does not readily infect Syrian hamsters. Antiviral Res 198, 105253 (2022).

33. Obeid, M., et al. Humoral Responses Against Variants of Concern by COVID-19 mRNA Vaccines in Immunocompromised Patients. JAMA Oncol (2022).

34. Mohammed, A.H., Blebil, A., Dujaili, J. & Rasool-Hassan, B.A. The Risk and Impact of COVID-19 Pandemic on Immunosuppressed Patients: Cancer, HIV, and Solid Organ Transplant Recipients. AIDS Rev 22, 151–157 (2020).

35. Fenwick, C., et al. Changes in SARS-CoV-2 Spike versus Nucleoprotein Antibody Responses Impact the Estimates of Infections in Population-Based Seroprevalence Studies. J Virol 95 (2021).

36. Maisonnasse, P., et al. COVA1-18 neutralizing antibody protects against SARS-CoV-2 in three preclinical models. Nat Commun 12, 6097 (2021).

37. Richardson, S.I., et al. HIV Broadly Neutralizing Antibodies Expressed as IgG3 Preserve Neutralization Potency and Show Improved Fc Effector Function. Front Immunol 12, 733958 (2021).

38. Punjani, A., Rubinstein, J.L., Fleet, D.J. & Brubaker, M.A. cryoSPARC: algorithms for rapid unsupervised cryo-EM structure determination. Nat Methods 14, 290–296 (2017).

39. Rubinstein, J.L. & Brubaker, M.A. Alignment of cryo-EM movies of individual particles by optimization of image translations. J Struct Biol 192, 188–195 (2015).

40. Pettersen, E.F., et al. UCSF ChimeraX: Structure visualization for researchers, educators, and developers. Protein Sci 30, 70–82 (2021).

41. Emsley, P., Lohkamp, B., Scott, W.G. & Cowtan, K. Features and development of Coot. Acta Crystallogr D Biol Crystallogr 66, 486–501 (2010).

42. Liebschner, D., et al. Macromolecular structure determination using X-rays, neutrons and electrons: recent developments in Phenix. Acta Crystallogr D Struct Biol 75, 861–877 (2019).

